# Retinal ganglion cell input to superior colliculus encodes salient information

**DOI:** 10.1101/2025.11.06.687035

**Authors:** Katharine Borges, Qiqi Xian, Rongwen Lu, Yajie Liang, Shannon Haley, Shirali Nigam, Na Ji

## Abstract

The superior colliculus (SC) is critical in detecting and responding to salient stimuli. However, it is unknown whether salience signals are inherited from external sources such as retina and cortex, or arise from the intrinsic circuitry within SC. We used in vivo 2-photon calcium imaging with adaptive optics to examine the activity of retinal ganglion cell (RGC) axon terminals in superficial SC (sSC). We found that RGC boutons exhibit similar saliency-related activity as sSC neurons. A majority of boutons respond more strongly to salient discontinuities of visual features, and change their orientation preference depending on stimulus properties such as the orientations of edges of drifting gratings. We furthermore demonstrate that these effects arise from the center-surround receptive field structure of RGCs. Our results show that the orientation tuning of RGCs is highly flexible, and that saliency encoding originates in retina rather than sSC.

## Introduction

The superior colliculus (SC) is critical in directing attention toward salient environmental stimuli^1–7^. Neurons in the superficial layer of the SC (sSC), which receives inputs from retina and visual cortex^8,9^, respond strongly to conspicuous visual features such as looming, contrasting motion, and cross-oriented patterns^10–13^. This enhanced activity encodes bottom-up saliency, which arises from physical features of stimuli that “pop out” of the visual scene and capture attention in an automatic, involuntary manner. Recently, we demonstrated that responses of sSC neurons are heightened in the retinotopic vicinity of salient discontinuities in the visual field (i.e. edges between a drifting grating and darkness, or between orthogonally or oppositely moving gratings), hence forming an activity map encoding salience in visual space^11^ as proposed in computational models of visual saliency^14,15^. Interestingly, grating edges also induce orientation tuning in sSC neurons, with their preferred orientation perpendicular to the edge orientation^11^.

In primates, saliency encoding is thought to arise in cortex or sSC^16–20^. In rodents, however, it remains unknown whether the saliency-related activity observed in sSC neurons is generated within sSC or inherited from the retinal ganglion cells (RGCs) or visual cortex. RGCs provide the main driving input for sSC neurons in mice; the sSC derives its retinotopic map, as well as orientation and direction tuning, from RGC activity^21–24^. In contrast, cortical inputs are believed to play a modulatory role in mouse sSC, enhancing the response of sSC neurons to looming objects^25^, suppressing responses to orthogonal gratings^26^, and mediating freezing behavior triggered by light flash^27^. Furthermore, previous work found no evidence of a saliency map in mouse primary visual cortex (V1) and demonstrated that responses of sSC neurons to salient visual stimuli were preserved in the absence of V1^11^. These results suggest that the encoding of salience in the mouse brain originates either in sSC itself, or from RGC inputs.

While retinal activity has not traditionally been considered to encode visual saliency, here we explicitly test whether RGCs underlie the responses to salient stimuli observed in the mouse sSC. Using in vivo two-photon calcium imaging of RGC axonal boutons in sSC, we find that a large majority of RGC boutons increase their activity in response to salient stimuli such as edges, oppositely moving gratings, and orthogonally moving gratings. Furthermore, oriented salient stimuli such as edges of drifting gratings induce orientation selectivity in RGC boutons, with their preferred orientation being perpendicular to the orientation of the edge. We find that these effects can arise from the center-surround receptive field structure of RGCs. Together, these results suggest that saliency encoding in the mouse originates in retina rather than SC or V1.

## Results

### In vivo two-photon imaging of RGC boutons in sSC reveals selectivity for orientation and direction with large-aperture drifting grating stimulation

To image RGC boutons in vivo, we injected adeno-associated virus encoding the calcium indicator GCaMP6s^28^ (AAV2/2.hSyn.GCaMP6s or AAV2/2.hSyn.GAP43.GCaMP6s) into the vitreous of the right eye of wildtype mice and implanted a cranial window over left SC. Overlying visual cortex was aspirated and the left lateral sinus was severed to gain access to a larger area of SC. Mice expressed GCaMP6s strongly throughout the retina (**Fig. 1a**), with dense RGC axonal projections in sSC (**Fig. 1b**).

**Figure 1.**
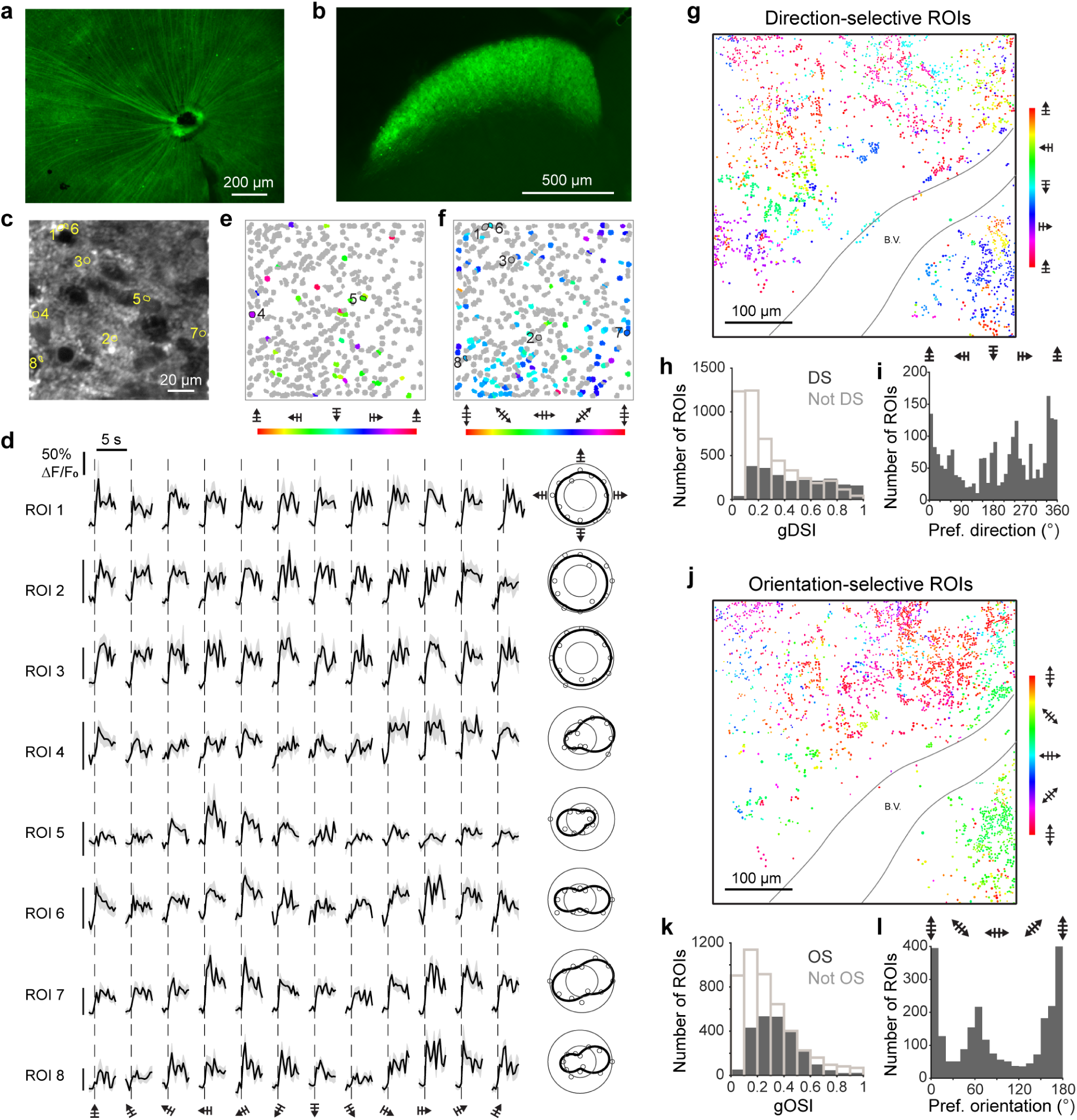
In vivo 2P calcium imaging of RGC boutons in sSC responding to large drifting gratings. (**a**) Retina flatmount from a mouse injected with AAV2/2-hSyn-GAP43-GCaMP6s into the vitreous of the eye. (**b**) Coronal brain section from the same mouse showing SC with fluorescent RGC axon terminals. (**c**) In vivo 2P fluorescence image of RGC boutons in sSC. (**d**) Example ΔF/F_0_ traces and polar plots for 8 regions of interest (ROIs) circled in **c**, averaged across stimulus repeats. Dashed line: onset of drifting grating. Shaded areas: ± S.E.M. Polar plots: outer circle represents maximal ΔF/F_0_; inner circle represents 50% of max ΔF/F_0_. (**e**) Direction-selective (DS) boutons color-coded by preferred direction. Gray boutons are visually responsive (max ΔF/F_0_ > 10%) but not DS. Bars indicate the grating orientation and arrows indicate direction of motion, respectively. (**f**) Orientation-selective (OS) boutons color-coded by preferred orientation. Gray boutons are visually responsive but not OS. (**g**) DS boutons color-coded by preferred direction in 25 FOVs over 450 μm × 450 μm in sSC. B.V.: blood vessel. (**h**) gDSI distributions for DS (gray bars; 2268 boutons, median 0.43) or non-DS (hollow bars; 4742 boutons, median 0.19) boutons in the FOVs in **g**. *p* value: 8.63e-154. (**i**) Preferred direction distribution for DS boutons. (**j**) OS boutons color-coded by preferred orientation for the same FOVs shown in **g**. (**k**) gOSI distributions for OS (gray bars; 2374 boutons, median 0.33) or non-OS (hollow bars; 4636 boutons, median 0.22) boutons in **j**. *p* value: 9.89e-80. (**l**) Preferred orientation distribution for OS boutons.

For in vivo imaging, mice were head-fixed with light anesthesia (1% isoflurane) and placed in a two-photon (2P) microscope with adaptive optics (AO) to minimize optical aberrations in the excitation light path^29,30^. As shown in a representative field of view (FOV) (**Fig. 1c**, 90 μm × 90 μm, 0.3 μm/pixel, 70 μm below the sSC surface), 2P fluorescent images revealed densely distributed putative RGC boutons. Regions of interest (ROIs) were either manually drawn around presumptive boutons or identified by Suite2p with manual curation^31^. Both methods of ROI segmentation yielded similar results.

We measured retinotopy to identify sSC areas that were mapped to specific regions of the visual stimulation screen, then recorded calcium transients from FOVs corresponding to the center of the stimulation screen. Large-aperture (80° visual angle) drifting gratings centered on the screen evoked robust activity in many boutons. We quantified the baseline-normalized change in GCaMP6s fluorescence (ΔF/F_0_) of each bouton as they responded to gratings drifting in one of twelve directions (from 0° to 330°, in 30° increments; example ROI 1-8, static gratings are presented for baseline; dashed lines indicate the onset of grating motion, **Fig. 1d**; Methods). A bouton was defined as visually responsive if its maximum ΔF/F_0_ evoked by at least one visual stimulus was more than 10%. Orientation- and direction-selectivity of visually responsive boutons were assessed by calculating the global orientation selectivity index (gOSI) and global direction selective index (gDSI), respectively, and testing for statistical significance with a permutation test with data shuffling^32^.

The majority of boutons in this FOV responded with similar strength to each of the grating directions and hence were untuned to direction or orientation (**Fig. 1d**, ROIs 1-3; **Fig. 1e,f**, gray ROIs). However, some boutons were direction-selective (DS; **Fig. 1d**, ROIs 4-5; color-coded by preferred direction, **Fig. 1e**), and some were orientation selective with similar responses towards gratings having the same orientation but moving in opposite directions (OS; **Fig. 1d**, ROIs 6-8; color-coded by preferred orientation, **Fig. 1f**), consistent with previous reports of DS and OS RGCs in the mouse retina^21,32,33^.

To examine orientation and direction tuning over a larger area of sSC, we imaged contiguous FOVs in another mouse (450 μm × 450 μm, acquired by tiling 25 90 μm × 90 μm FOVs, 0.3 μm/pixel, 50 μm below surface) and characterized the orientation and direction selectivity of individual boutons (**Fig. 1g-l**). 47% of 7010 visually responsive boutons were found to be untuned, while 32% and 34% were DS and OS (color-coded by preferred direction or orientation, **Fig. 1g,j**), respectively.

As expected, distributions of gDSI showed that DS boutons tended to have higher gDSI values than visually responsive but non-DS boutons, with significantly different medians (**Fig. 1h**, **Supp. Table 1**; Kolmogorov-Smirnov test was used to compare cumulative distributions for all figures; see supplementary tables for all medians and p values). Although all directions were represented, the distribution for preferred direction showed a bias for upward and posterior motion (**Fig. 1i**). OS boutons tended to have higher gOSI values than visually responsive but not OS boutons, again as expected (**Fig. 1k**; **Supp. Table 1**). The distribution for preferred orientation was dominated by horizontal and near-vertical gratings (**Fig. 1l**).

The activity of nearby boutons was more strongly correlated than that of distant boutons, in accordance with prior results^21^, and boutons with similar orientation and direction preference were clustered together (**Fig. 1g,j**; zoomed-in views, **Supp. Fig. 1a,b**). Both activity correlation and direction/orientation clustering extended over tens of µm (**Supp. Fig. 1c,d**), consistent with anatomical measurements of RGC axon arborization in the sSC^34^. Together, these observations suggest that boutons within each cluster may originate from the same RGC, or RGCs with similar tuning properties.

### Edge of circular gratings induces orientation tuning and increases calcium activity of RGC boutons

Previous work in our lab showed that neurons in sSC respond more strongly to the edge of circular gratings^11^. This finding is consistent with the ideas put forth by Itti and Koch in their computational model of visual saliency; i.e. discontinuities of features within the visual field – in our case, the sharp boundary between grating and darkness – elicit higher responses from neurons that are retinotopically mapped to these salient features^14,15^. In addition, we observed that stimulus edges induce orientation tuning in sSC neurons, with the preferred orientation perpendicular to the orientation of the edge^11^. Given that we did not observe this in V1 activity^11^, we sought to determine whether these properties exist in retinal input.

We first mapped the retinotopy of RGC boutons to find the region of sSC corresponding to the middle of the stimulation screen. We then acquired 2P fluorescence images while presenting either a large-aperture (80° visual angle, **Fig. 2a**) or small-aperture (18.5° visual angle, **Fig. 2b**) drifting grating, with the small grating positioned such that the edge of the grating fell within the retinotopic area of sSC that was being imaged (green or orange boxes, **Fig. 2a,b**).

**Figure 2.**
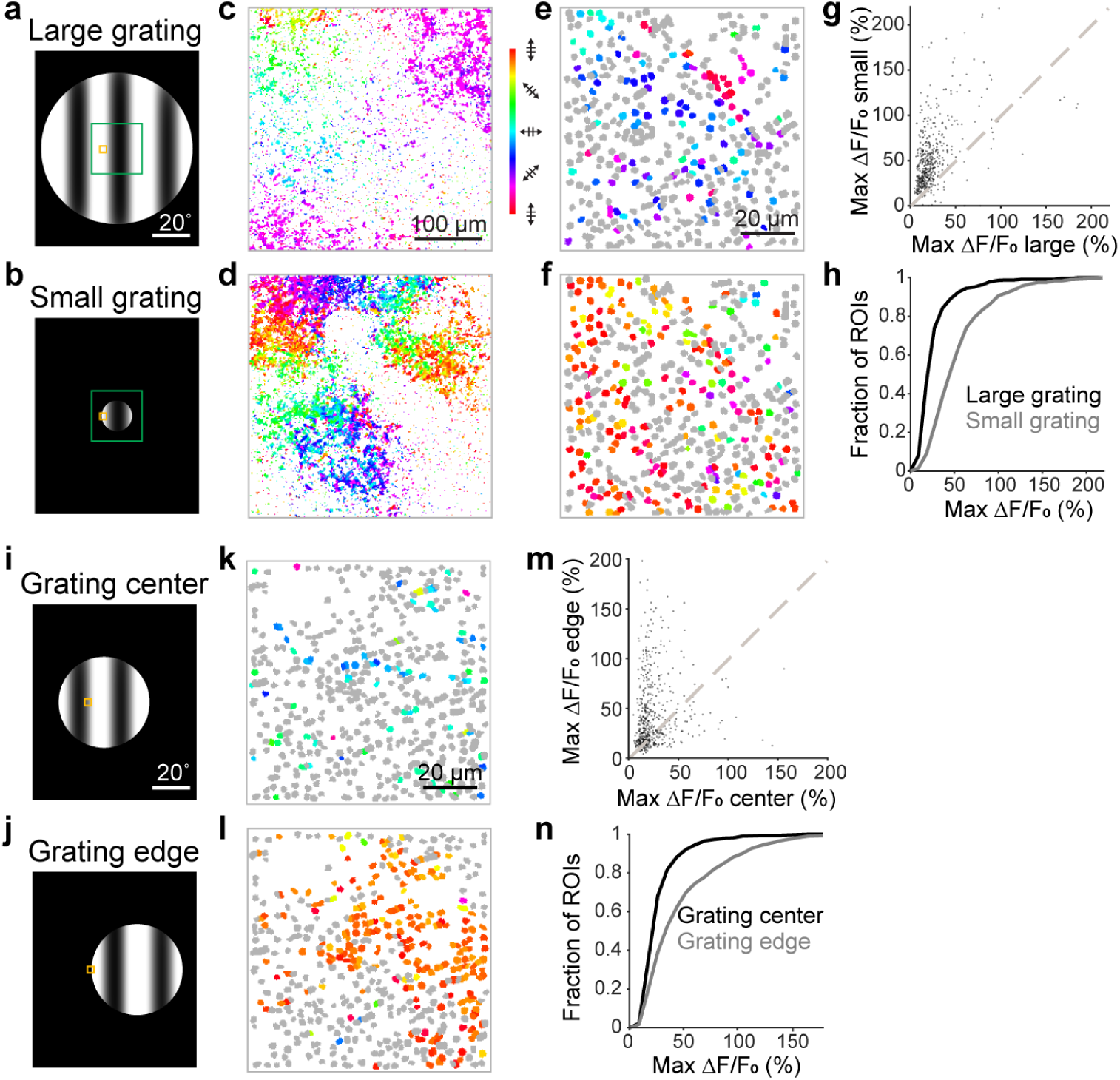
Edge of circular gratings induces orientation selectivity and increases calcium activity of RGC boutons. (**a**) Large- and (**b**) small-aperture drifting gratings. Green and orange boxes: retinotopic area of FOVs for **c,d** and **e,f**, respectively. (**c**) OS pixels in an example FOV (pixel size: 1.2 μm) color-coded by preferred orientation measured with large gratings. (**d**) Same FOV as in **c**, orientation preference measured with small gratings. (**e**) RGC boutons in an example FOV (pixel size: 0.3 μm) color-coded based on preferred orientation measured with large gratings. Gray: non-OS boutons; colored: OS boutons. (**f**) Same FOV as **e** but now stimulated by the edge of small gratings. (**g**) Scatter plot of max ΔF/F_0_ of each bouton for small versus large gratings, and (**h**) cumulative distributions of max ΔF/F_0_ for large (black) or small (gray) gratings (medians: 19.2% and 44.5%, respectively). *p* value: 2.46e-59. (**i,j**) Mid-sized drifting gratings with imaged FOV (orange boxes) mapped to center or edge of a mid-size grating, respectively. (**k,l**) RGC boutons from an example FOV color-coded based on preferred orientation from grating-center or grating-edge stimulation, respectively. Gray: non-OS boutons; colored: OS boutons. (**m**) Scatter plot of max ΔF/F_0_ of each bouton for grating edge vs. center, and (**n**) cumulative distributions of max ΔF/F_0_ for grating center (black) and edge (gray) of the grating (medians: 19.1% and 30.4%, respectively). *p* value: 1.60e-21.

Orientation analysis of image pixels within a 360 μm × 360 μm FOV (green box, **Fig. 2a,b**; 1.2 μm/pixel, **Fig. 2c**) revealed sparse and clustered OS pixels measured with large gratings, consistent with **Fig. 1**. But with the small grating stimuli, a much higher proportion of pixels became OS at the edge of small gratings (**Fig. 2d**), suggesting that the grating edge induced changes in orientation tuning of RGC boutons, similar to what we previously observed in sSC neurons^11^.

To assess the orientation selectivity and response magnitude of individual RGC boutons retinotopically mapped to the edge of small gratings, we imaged 90 μm × 90 μm FOVs at higher resolution (orange box, **Fig. 2a,b**; 0.3 μm/pixel). For the same bouton population, the presence of the edge of small gratings increased the fraction of OS boutons from 29% to 44% and induced a drastic change in preferred orientation of many OS boutons in a representative FOV (**Fig. 2e,f**).

The grating edge also increased the calcium response of the vast majority of RGC boutons, with maximum ΔF/F_0_ values significantly higher for the small grating edge than for the large grating (**Fig. 2g,h**; 90% of boutons above unity line; **Supp. Table 2**). The same trend held across population data (**Supp. Fig. 2a-d; Supp. Table 3**; 6943 visually responsive boutons from 13 FOVs of 8 mice).

To ensure that the observed trends did not result simply from the difference in the overall size of the grating, we used a mid-size grating (52° of visual angle) and moved its location so that either the middle or edge of the grating covered the retinotopic area for a given FOV (orange box, **Fig. 2i,j**). More boutons were orientation tuned and had higher responses at the edge of the grating compared to the middle (example FOV, **Fig. 2k-n**; population data, 1125 boutons from 2 FOVs of 2 mice, **Supp. Fig. 2e-h**, **Supp. Table 2,3**), indicating that the presence of the edge, rather than the size of the grating, produced the observed changes in orientation selectivity and max ΔF/F_0_.

Similar to what we previously observed in sSC neurons^11^, orientation selectivity was induced by the edge of gratings in RGC boutons down to depths of ∼200 μm below sSC surface, with patches of similar preferred orientation persisting for tens of microns (**Supp. Fig. 3**). Edge-induced tuning was also present across various spatial and temporal frequencies of the grating (**Supp. Fig. 4**).

### RGC boutons change their tuning to prefer orientations that are perpendicular to straight edges

Previously, edges of grating stimuli were found to induce sSC neurons to prefer the grating orientation that was perpendicular to the edge^11^. To test this in retinal inputs, we generated a vertical or horizontal edge by superimposing a dark rectangle over large drifting gratings, and imaged RGC boutons retinotopically mapped to the edge (**Fig. 3a**). Pixel-based analysis of a 450 μm × 450 μm FOV (green box, **Fig. 3a**; 1.5 μm/pixel, 40 μm below the sSC surface, **Fig. 3b**) revealed a much higher proportion of OS pixels in the presence of an edge with the predominant orientation preference being perpendicular to the edge (**Fig. 3c**).

**Figure 3.**
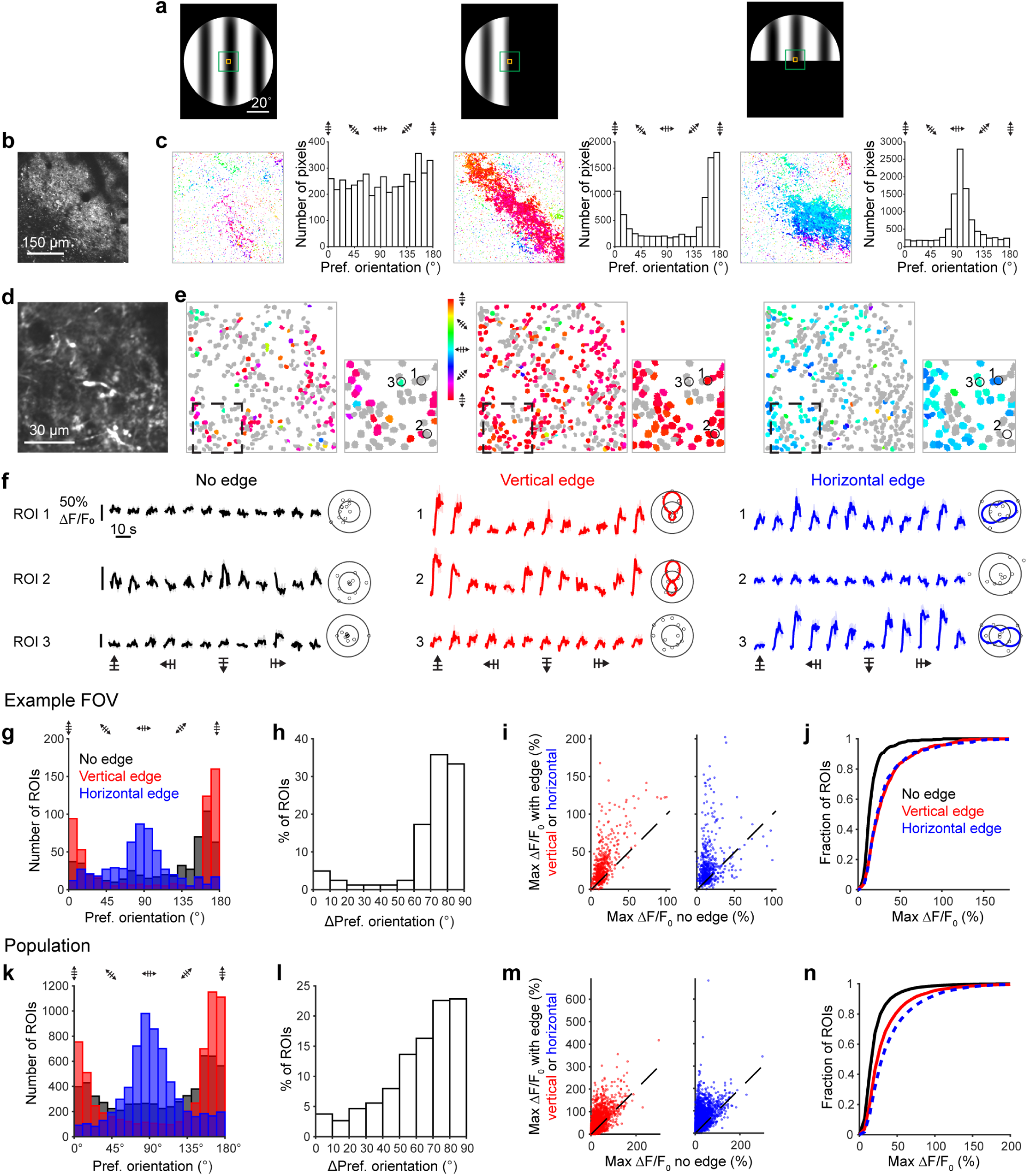
RGC boutons change their tuning to prefer grating orientations that are perpendicular to straight edges. (**a**) Stimuli: large drifting grating (80° visual angle) with either no edge or with a straight vertical or horizontal edge. Green and orange boxes: retinotopic area of FOVs in **b**, **d**, respectively. (**b**) 2P image of an example 450 μm × 450 μm FOV. Pixel size: 1.5 μm. (**c**) OS pixels color-coded by preferred orientation (left) and preferred orientation distributions for pixels (right) in **b** for each stimulus in **a**. (**d**) Image of RGC boutons in an example 90 μm × 90 μm FOV. Pixel size: 0.3 μm. (**e**) RGC boutons from **d**, color-coded based on orientation selectivity in response to stimuli in **a**. Gray: non-OS boutons; colored: OS boutons. Insets: Enlarged views of boutons in dashed boxes. (**f**) ΔF/F_0_ traces and polar plots for the three circled ROIs in **e**, responding to gratings with no edge (black), vertical edge (red), and horizontal edge (blue). Shaded areas: ± S.E.M. (**g**) Preferred orientation distributions for boutons in **d** that are OS for gratings with no edge (black), vertical edge (red), and horizontal edge (blue). (**h**) Change in preferred orientation between vertical and horizontal edges, for boutons in **d** that are OS with both edges. (**i**) Scatter plots of max ΔF/F_0_ for 527 visually responsive boutons in **d** with vertical (left) or horizontal (right) edge vs. no edge. 85% and 75% of boutons above unity line for vertical and horizontal edge, respectively. (**j**) Cumulative distributions of max ΔF/F_0_ for visually responsive boutons in **d** for each stimulus (medians for no edge, vertical edge, and horizontal edge: 13.2%, 24.0%, 22.7%, respectively). *p* values: 6.57e-33 for no edge vs. vertical edge, 5.22e-30, for no edge vs. horizontal edge, 0.64 for vertical vs. horizontal edge. (**k-n**) Same as **g-j**, for 6125 boutons in 11 FOVs from 5 mice. **m**: 76% and 82% of boutons above unity line for vertical and horizontal edges, respectively. **n**: Medians for no edge, vertical edge, and horizontal edge: 15.5%, 23.5%, 29.7%, respectively. *p* values: 6.37e-167 for no edge vs. vertical edge, 0 for no edge vs. horizontal edge, 3.64e-40 for vertical vs. horizontal edge.

Using high-resolution imaging, we investigated how individual RGC boutons modified their tuning properties in response to straight edges. In an example 90 μm × 90 μm FOV (orange box, **Fig. 3a**; 0.3 μm/pixel, **Fig. 3d**), the presence of an edge increased the number of OS boutons (**Fig. 3e**, **Supp. Table 4**). When a vertical edge was present, most OS boutons preferred horizontally oriented gratings; conversely, when a horizontal edge was present, most OS boutons preferred vertical gratings. Some boutons remained tuned with both edges, changing their preferred orientation to be perpendicular to the presented edge (ROI 1, **Fig. 3f**), while other boutons were only OS for one of the edges (ROI 2 and 3, **Fig. 3f**).

Within this FOV, boutons that were OS under each stimulus condition showed drastic changes in their preferred orientation distributions (**Fig. 3g**). Strikingly, many boutons changed their orientation preference by 45° or more with different edge orientations: out of 81 boutons that were tuned to both edges, 73 (90%) changed orientation preference by ≥ 45° (**Fig. 3h**). Consistent with our observations at the edge of circular gratings (**Fig. 2**), we found that max ΔF/F_0_ was significantly higher in the presence of an edge for the vast majority of visually-responsive boutons in this FOV (**Fig. 3i,j**; **Supp. Table 4**).

We observed similar trends across multiple FOVs and mice (6125 boutons responsive to at least one stimulus, 11 FOVs, 5 mice; **Fig. 3k-n**, **Supp. Table 4**). Preferred orientation distributions were different for each edge, with peak orientations orthogonal to the respective edges (**Fig. 3k**); 80% of the 932 boutons that were tuned for both edges changed their orientation preference by ≥ 45° (**Fig. 3l**); and a large majority of boutons increased max ΔF/F_0_ when an edge was present (**Fig. 3m,n**). We observed similar trends for 45° and 135° oblique edges across multiple FOVs and mice (2192 boutons, 4 FOVs in 3 mice; Supp. Fig. 5, Supp. Table 5).

### Presence of an edge modifies the orientation tuning properties of both natively tuned and non-natively tuned RGC boutons

While we found that the presence of an edge increased the density of tuned RGC boutons, a minority of boutons were OS with large gratings alone, consistent with prior studies of RGC cell bodies and axon terminals that utilized gratings or bars extending over a large portion of the visual field^21,32,33^. Having shown that the presence of an edge can manipulate the orientation selectivity of RGC boutons, we next sought to determine whether natively tuned boutons (i.e., those that are OS in the presence of a large grating, which we define as the “native” condition) behave differently in the presence of an edge than non-natively tuned boutons (i.e., those that were not tuned with a large grating, but became tuned when a horizontal or vertical edge was present). Given that the intrinsic orientation tuning of some RGCs arises from their unique dendritic morphology and presynaptic inputs^33^, we investigated whether natively tuned RGC boutons would be more resistant to changing their orientation selectivity than non-natively tuned boutons.

Across the population (same as in **Fig. 3k-n**), 1128 boutons were natively tuned and predominantly preferred horizontal gratings moving up or down (**Fig. 4a**). 67% of these remained tuned in the presence of the vertical edge, which favors tuning for horizontal gratings and hence matched the predominant native tuning (**Fig 4b**, red). However, when a horizontal edge (which favors tuning for vertical gratings) was present, most (66%) boutons that were natively tuned for horizontal gratings became untuned. Less than half of all natively tuned boutons (43%) remained tuned with a horizontal edge, and most of these changed their tuning to prefer vertical gratings moving left or right (**Fig. 4b**, blue). For 2878 non-natively tuned boutons, 64% and 65% became tuned in the presence of the vertical or horizontal edge, respectively, with the presence of an edge eliciting similar orientation distributions to those of natively tuned boutons (**Fig. 4h**).

**Figure 4.**
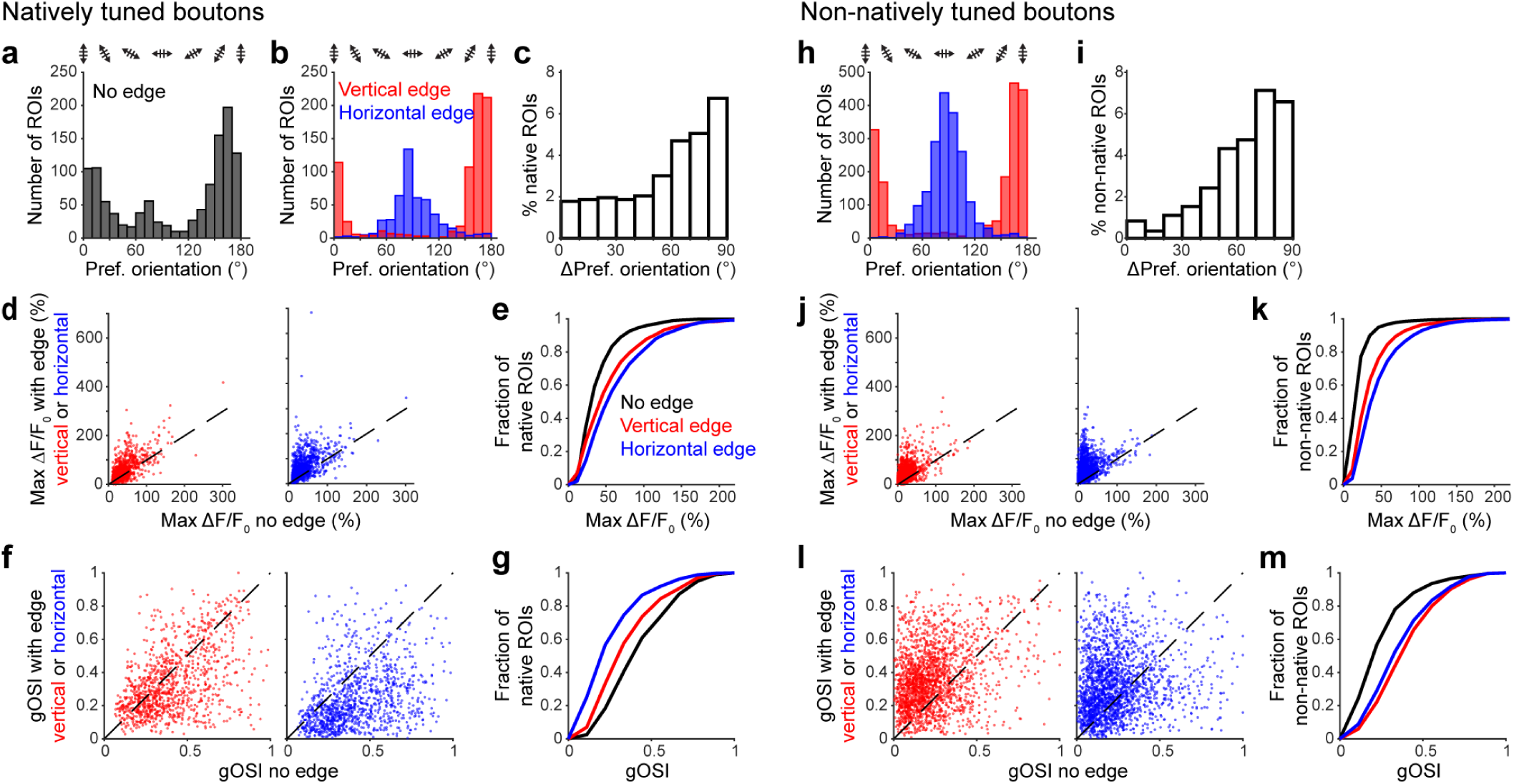
Presence of an edge modifies the orientation tuning properties of both natively tuned and non-natively tuned RGC boutons. Data are from the same boutons as Fig. 3k**-n**. (**a**) Preferred orientation distribution for natively tuned boutons with large grating (no edge). (**b**) Preferred orientation distributions for boutons that are OS for gratings with vertical (red) or horizontal (blue) edge. (**c**) Change in preferred orientation between vertical and horizontal edges, for natively tuned boutons that are OS with both edges. (**d**) Scatter plots of max ΔF/F_0_ for natively tuned boutons with vertical (left) or horizontal (right) edge versus no edge. 68% and 78% of natively tuned boutons above unity line for vertical and horizontal edges, respectively. (**e**) Cumulative distributions of max ΔF/F_0_ for natively tuned boutons for each stimulus. Medians for no edge, vertical edge, and horizontal edge: 30.5%, 41.9%, 50.9%, respectively. *p* values: 2.95e-19 for no edge vs. vertical edge, 1.30e-44 for no edge vs. horizontal edge, 7.83e-9 for vertical vs. horizontal edge. (**f**) Scatter plots of gOSI for natively tuned boutons with vertical (left) or horizontal (right) edge vs. no edge. 39% and 19% of boutons above unity line for vertical and horizontal edges, respectively (**g**) Cumulative distributions of gOSI for natively tuned boutons. Medians for no edge, vertical edge, and horizontal edge: 0.38, 0.29, 0.20, respectively. *p* values: 3.28e-17 for no edge vs. vertical edge, 3.61e-78 for no edge vs. horizontal edge, 5.23e-30 for vertical vs. horizontal edge. (**h-m**) Same as **b-g**, for non-natively tuned boutons. **j**: 83% and 90% of boutons above unity line for vertical and horizontal edges, respectively. **k**: Medians for no edge, vertical edge, and horizontal edge: 14.6%, 27.6%, 36.4%, respectively. *p* values: 5.60e-186 for no edge vs. vertical edge, 0 for no edge vs. horizontal edge, 3.02e-33 for vertical vs. horizontal edge. **l**: 76% and 65% of boutons above unity line for vertical and horizontal edges, respectively. **m**: Medians for no edge, vertical edge, and horizontal edge: 0.20, 0.36, 0.32, respectively. *p* values: 1.18e-137 for no edge vs. vertical edge, 1.65e-84 for no edge vs. horizontal edge, 1.04e-10 for vertical vs. horizontal edge.

The presence of an edge had similar effects on natively and non-natively tuned boutons. Among the natively tuned boutons that stayed OS with both edges, 72% changed their preferred orientation by at least 45° between the vertical and horizontal edge (**Fig. 4c**). For non-natively tuned boutons that were OS with both edges, 83% changed their preferred orientation by ≥ 45° (**Fig. 4i**). Hence, a similar proportion of natively tuned and non-natively tuned boutons showed a large change in their preferred orientation between the two edges. In addition, a majority of both natively and non-natively tuned boutons had higher max ΔF/F_0_ with either edge (**Fig. 4d,e,j,k**; 68% and 78% of natively tuned boutons above unity line for vertical and horizontal edges, respectively; for non-natively tuned boutons these percentages were 83% and 90%), with the medians of max ΔF/F_0_ distributions for edges significantly higher than for the large grating (**Supp. Table 6**).

A striking difference between the natively tuned and non-natively tuned populations was how their strength of tuning changed in the presence of an edge. Natively tuned boutons showed a significant drop in gOSI for either edge, indicating weaker tuning (**Fig. 4f,g; Supp. Table 6**). In contrast, gOSI increased significantly for non-natively tuned boutons that became tuned with either edge, as expected since they are untuned in the no-edge condition (**Fig. 4l,m; Supp. Table 6**; see **Supp. Fig. 6,7** for further analysis of non-natively tuned boutons). Similar but somewhat weaker effects were observed for oblique edges (**Supp. Fig. 6,7; Supp. Table 7,8**).

Taken together, these results suggest that the preferred orientation of both natively tuned and non-natively tuned RGC boutons can be manipulated by the presence of an edge. However, the circuit mechanisms that underlie tuning of natively OS boutons can partially counteract the edge-induced effect such that boutons become more weakly tuned if forced away from their natively preferred orientation.

### Model of center-surround receptive field structure reproduces experimentally observed edge effects

We hypothesized that the center-surround organization of RGC receptive fields (RFs) underlies the enhancement of activity and the change in orientation tuning observed at edges. To test this, we used a 2D Difference-Of-Gaussians (DOG) function to model a circularly symmetric RF^35^. The excitatory center and inhibitory surround were represented by two Gaussian functions with amplitudes of 1.8 and 1 and RMS widths of 22.4 μm and 30.0 μm^36–38^, respectively (cross section line profiles, **Fig. 5a** teal and purple curves; Methods), resulting in a DOG function that exhibited a “Mexican-hat” profile with a central peak surrounded by a negative trough (**Fig. 5a** gray curve; **Fig. 5b** 3D view).

**Figure 5:**
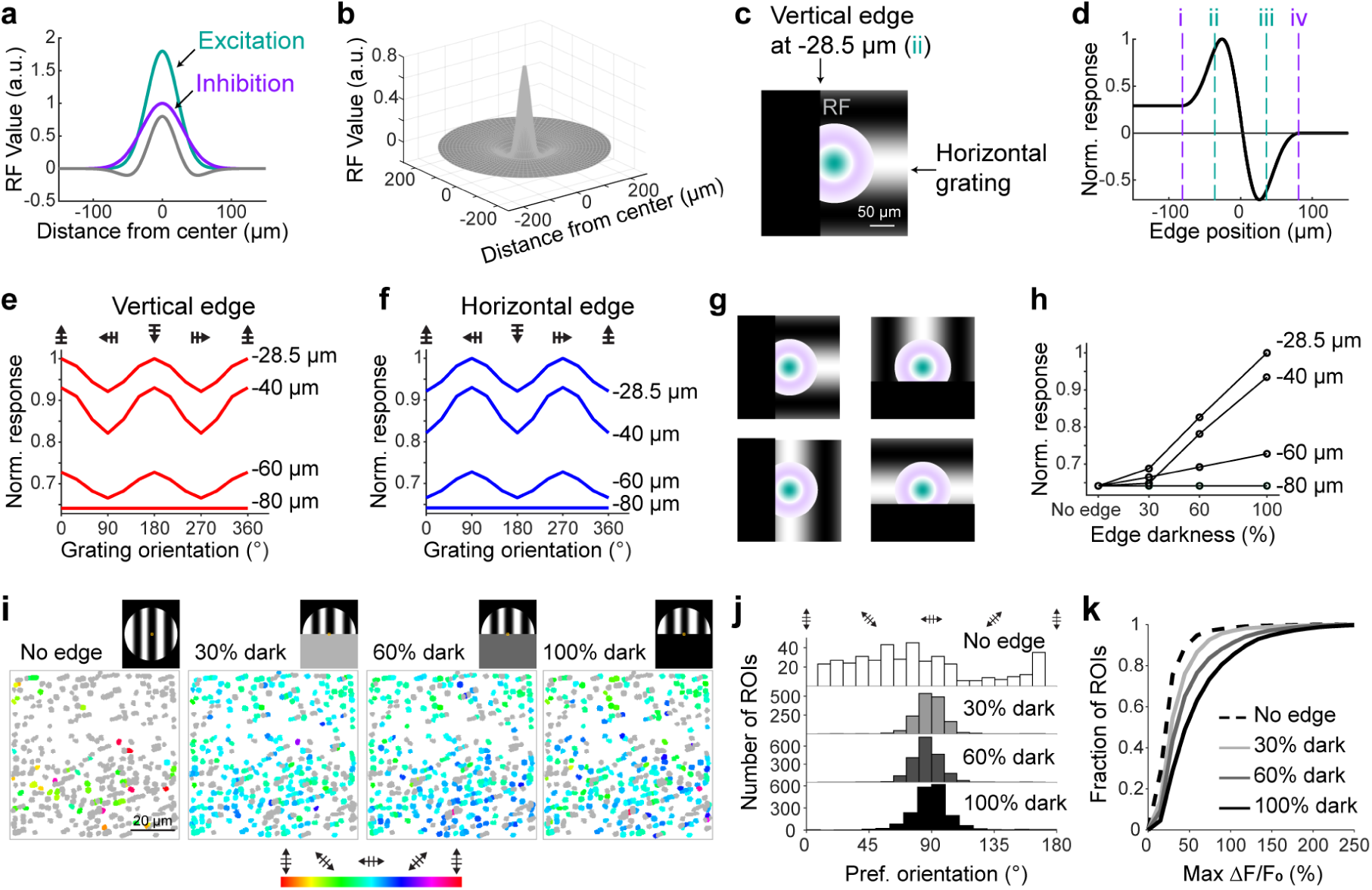
Difference-Of-Gaussians simulation of RGC receptive field yields similar results to experiments. (**a**) Gaussian curves for the excitatory and inhibitory regions (teal and purple curves), and net (gray curve, excitation minus inhibition). (**b**) 3D Plot of the RF. (**c**) Top-down view of the RF showing the excitatory center (teal) and inhibitory surround (purple), and the position of the vertical edge of the horizontal grating relative to the RF that gives the highest response (28.5 μm to the left of the RF center). (**d**) Normalized responses of the RF with a horizontal grating and vertical edge, for different edge positions relative to the center of the RF (at 0 μm). Vertical dashed lines: (i) left edge of inhibitory surround, (ii) border of inhibitory surround and excitatory center, (iii) border of excitatory center and inhibitory surround, (iv) right edge of inhibitory surround. (**e**) Normalized response of RF to gratings of different orientations, with a vertical edge at different positions relative to the RF center. (**f**) Same as **e**, for a horizontal edge. (**g**) Schematics of a vertical edge (left) and horizontal edge (right) with gratings orthogonal or parallel to an edge positioned at −40 μm. (**h**) Normalized response of the RF to a horizontal grating with a vertical edge of different gray levels and edge positions. **i**) Boutons in an example 76.8 μm × 76.8 μm FOV, color-coded based on orientation selectivity in response to a large drifting grating (80° visual angle) presented with either no edge or with a horizontal edge of 30%, 60%, or 100% darkness. Gray boutons: visually responsive but not OS; colored boutons: OS. (**j**) Preferred orientation distributions for boutons that are OS with each stimulus. (**k**) Cumulative distributions of max ΔF/F_0_ for 3966 boutons in 7 FOVs from 4 mice, for each stimulus. Medians for no edge, light gray, dark gray, and black edge: 19.9%, 27.6%, 33.3%, 45.5%, respectively. *p* values: 2.99e-87 for no edge vs. light gray edge, 8.57e-187 for no edge vs. dark gray edge, 0 for no edge vs. black edge, 4.98e-24 for light gray edge vs. dark gray edge, 8.34e-138 for light gray edge vs. black edge, 2.90e-49 for dark gray edge vs. black edge.

We then calculated the responses evoked by a horizontally oriented grating with a vertical edge (**Fig. 5c**) to determine how they varied as a function of the edge’s location within the RF (**Fig. 5d**). When the edge was far to the left of the RF, the grating covered the entire RF, yielding a weakly excitatory response (left edge of RF, −80.5 μm: dashed line i, **Fig. 5d**). As the edge shifted to the right, the inhibitory drive decreased and the response increased, reaching maximum when the edge was near the inhibition-excitation border (−28.5 μm, vertical edge location in **Fig. 5c**; dashed line ii, **Fig. 5d**). As the edge moved further to the right, excitation decreased faster than inhibition and the response started to decrease, reaching a minimum near the excitation-inhibition border (dashed line iii, **Fig. 5d**). Finally, the response went to zero when the grating moved beyond the right border of the inhibitory region and no longer overlapped with the RF (dashed line iv, **Fig. 5d**). This accords with our experimental results showing that RGCs with their RFs located near the edge had the strongest activity.

To test whether the center-surround RF enables flexible orientation tuning, we evaluated the response evoked by drifting gratings of different orientations with either a vertical or a horizontal edge. With the vertical edge at the position that resulted in maximum response (−28.5 μm relative to center), we found that horizontally oriented gratings evoked the largest responses (normalized to peak response, top trace, **Fig. 5e**). This preference held for a range of edge positions (bottom traces, **Fig. 5e**). Similarly, a horizontal edge resulted in a preference for vertically oriented gratings for a range of edge positions (**Fig. 5f**).

This change in orientation preference can be explained by the relative overlap between the grating and the inhibitory and excitatory portions of the RF. As shown for RFs with an edge located at −40 μm relative to center (**Fig. 5g**), the excitatory center has similar overlap with the grating whether the edge is vertical or horizontal. However, a smaller portion of the inhibitory surround overlaps the grating when the edge is perpendicular (top row, **Fig. 5g**) than when the edge is parallel (bottom row, **Fig. 5g**) to the grating orientation. As a result, gratings that are orthogonal to the edge will stimulate less inhibition and elicit a higher response from the RF. Hence, consistent with our experimental results, the DOG RF led to a preference for horizontal gratings when a vertical edge was present and a preference for vertical gratings when a horizontal edge was present.

If the enhancement of RGC activity at edges results from the center-surround organization of their RFs, we would expect to see smaller max ΔF/F_0_ values for lighter edges, as lighter edges should increase lateral inhibition and reduce RGC activity. To this end, we simulated edges with different darkness levels and found that the maximum response of the RF increased monotonically with edge darkness, as expected (**Fig. 5h**). To test this experimentally, we presented animals with a large grating and either no edge, a light gray (30% darkness), dark gray (60% darkness), or fully black (100% darkness) horizontal edge (**Fig. 5i-k**). As before, we imaged RGC boutons at high resolution from FOVs at the retinotopic position of the edge. We found a substantial increase in the fraction of OS boutons with a preference for vertically oriented gratings (example FOV, **Fig. 5i**; population result of 3966 visually responsive boutons in 7 FOVs from 4 mice, **Fig. 5j**; the percentages of OS boutons were 12%, 41%, 50% and 48% with no edge, light gray, dark gray, and black, respectively). Consistent with the DOG model, we observed a shift in the max ΔF/F_0_ distribution to significantly higher values with darker edges (**Fig. 5k**; **Supp. Table 9**).

In summary, our experimental results could be explained with a circularly symmetric center-surround RF model without invoking synaptic plasticity or other dynamic mechanisms. A similar model developed for sSC neurons also demonstrated that center-surround RFs could underlie a switch in preferred orientation with different edge orientations^39^, as shown in our earlier work in sSC neurons^11^.

### RGC bouton activity is enhanced and tuning modified at salient borders between orthogonally and oppositely moving gratings

The sSC responds to a diverse range of salient visual stimuli. In particular, neurons within the sSC have been shown to enhance their activity when presented with concentric gratings moving in opposite directions or oriented orthogonally to one another^10,13,26,40^. Our previous work also indicated that sSC neurons mapped to the border between the concentric gratings encode saliency with enhanced activity and induced orientation tuning^11^. We therefore investigated whether RGC boutons respond similarly to these stimuli.

We first imaged from a 450 μm × 450 μm FOV to localize the area responsive to a small (18.5° visual angle) drifting grating (green box, **Fig. 6a**; pixel size: 1.5 μm, **Fig. 6b**). We then presented a large uniform drifting grating (80° visual angle) or concentric gratings (orthogonally or oppositely moving gratings), with the inner grating at the same size and position as the small grating, and the outer grating annulus with the same outer diameter as the large grating (**Fig. 6a**). Pixel-based analysis of this FOV revealed more OS pixels in the presence of concentric gratings compared to the large uniform grating (**Fig. 6b**). Within the area retinotopically mapped to the inner grating (dashed circles, **Fig. 6b**), the preferred orientations of the pixels were strongly affected by the presence of an outer annulus, with dominant preferences being nearly 90° apart for the orthogonally and oppositely moving gratings (**Fig. 6c**).

**Figure 6.**
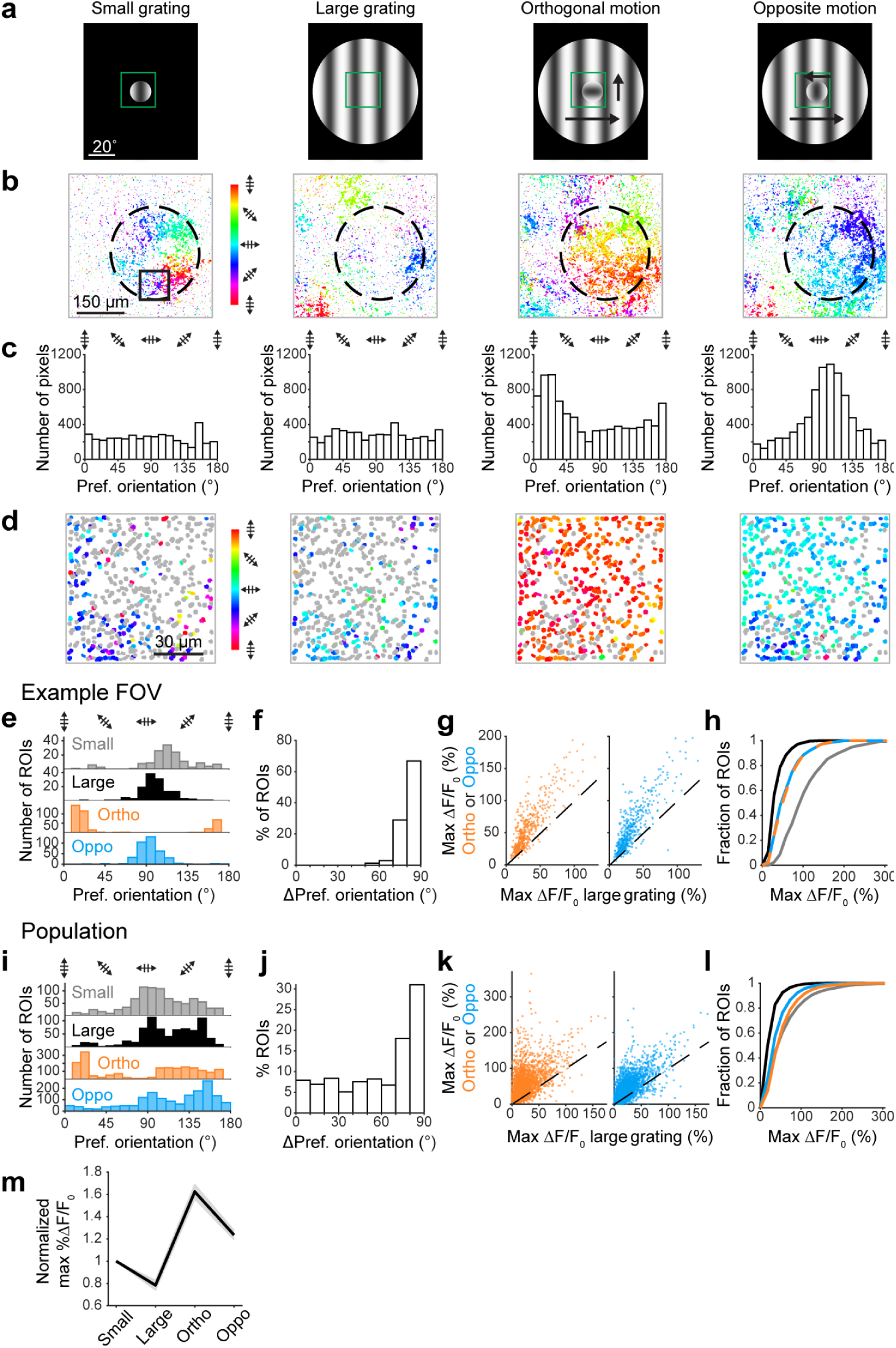
RGC boutons encode saliency of orthogonally and oppositely moving gratings. (**a**) Stimulus types: small (18.5° visual angle), large (80° visual angle), orthogonally moving, and oppositely moving gratings. (**b**) OS pixels in an example 450 μm × 450 μm FOV color-coded based on preferred orientation in response to stimuli in **a**. Pixel size: 1.5 μm. (**c**) Preferred orientation of pixels within the dashed circle in **b**, for each stimulus type. Note that for orthogonal gratings, the OS map refers to orientation of the inner grating. (**d**) Boutons in box shown in **b**, color-coded based on preferred orientation in response to stimuli in **a**. Gray: visually responsive but not OS boutons; colored: OS boutons. Pixel size: 0.3 μm. (**e**) Preferred orientation distributions for boutons that are tuned with each stimulus type within the FOV in **d**. (**f**) Change in preferred orientations between orthogonally and oppositely moving gratings, for boutons in **d** that are OS with both stimuli. (**g**) Scatter plots of max ΔF/F_0_ for boutons in **d** with oppositely (left) or orthogonally (right) moving gratings vs. large uniform gratings. 94% and 92% of boutons above unity line for orthogonally and oppositely moving gratings, respectively (**h**) Cumulative distributions of max ΔF/F_0_ for each stimulus type for boutons in **d**. Medians for large, small, orthogonal, and oppositely moving gratings: 26.5%, 91.0%, 45.8%, 49.9%, respectively. *p* values: 2.13e-106 for large vs. small grating, 2.97e-31 for large vs. ortho. gratings, 1.52e-25 for large vs. oppo. gratings, 1.52e-37 for small vs. ortho. gratings, 5.06e-39 for small vs. oppo. gratings, 0.272 for ortho. vs. oppo. gratings. (**i-l**) Same as **e-h**, for 3224 boutons in 6 FOVs from 4 mice. **k**: 89% and 84% of boutons above unity line, respectively. **l**: Medians for large, small, orthogonal, and oppositely moving gratings: 18.1%, 41.9%, 41.4%, 31.7%, respectively. *p* values: 1.33e-209 for large vs. small grating, 2.49e-276 for large vs. ortho. gratings, 1.85e-127 for large vs. oppo. gratings, 1.20e-16 for small vs. ortho. gratings, 2.01e-39 for small vs. oppo. gratings, 6.59e-33 for ortho. vs. oppo. gratings. (**m**) Max ΔF/F_0_ for each stimulus type, normalized to the max ΔF/F_0_ of each bouton for small gratings, averaged for all boutons in **i-l**.

We then imaged a 90 μm × 90 μm FOV corresponding to the edge of the inner grating at single-bouton resolution (box in **Fig. 6b**; pixel size: 0.3 μm, **Fig. 6d**). Consistent with our results at curved and straight edges between gratings and dark screen, the border between concentric gratings induced dramatic changes in preferred orientation and density of OS RGC boutons (**Fig. 6d,e**). In this FOV, 311 out of 311 (100%) boutons that were tuned with both orthogonally and oppositely moving gratings changed their preferred orientation by at least 45° (**Fig. 6f**). In addition, the max ΔF/F_0_ of most boutons was higher with concentric gratings than large uniform gratings (**Fig. 6g,h**), but not as high as for the edge of the small grating (**Fig. 6h; Supp. Table 10**). The same trends were observed from 3224 visually responsive boutons in 6 FOVs from 4 mice (**Fig. 6i-l; Supp. Table 10**).

As before, we split the boutons into populations that were natively tuned (OS with large uniform gratings) and non-natively tuned (OS with concentric but not large uniform gratings). Similar to our findings with straight edges (**Fig. 3**), the majority of both populations changed their preferred orientation and increased their max ΔF/F_0_ at the border between concentric gratings (**Supp. Fig. 8a-e,h-k; Supp. Table 11**). Out of 738 natively tuned boutons, 32% changed their preferred orientation by at least 45° between orthogonally and oppositely moving gratings (**Supp. Fig. 8c**); for the 1765 non-natively tuned boutons, this percentage was 29% (**Supp. Fig. 8i**). In addition, natively tuned boutons had a significantly lower gOSI with concentric gratings compared to large uniform gratings (**Supp. Fig. 8f,g; Supp. Table 11**), while gOSI values for non-natively tuned boutons were significantly higher (**Supp. Fig. 8l,m; Supp. Table 11**). (See **Supp. Fig. 9 and Supp. Table 12** for further analysis of non-natively tuned boutons.) This accords with our results for straight edges (**Fig. 4f,g,l,m**), suggesting that the same mechanisms drive these changes in tuning properties at the edge of a grating and the border between concentric gratings.

Lastly, we examined how RGC boutons’ response magnitudes varied with stimulus type (**Fig. 6m**). For every bouton, we normalized its max ΔF/F_0_ for each stimulus to its value for small gratings. We then averaged across boutons. Similar to our previous observations in sSC neurons^11^, RGC boutons responded more strongly to salient visual features such as the edge of the small grating or the border between the concentric gratings than to the large uniform grating. However, activity for orthogonally moving gratings was higher than for oppositely moving gratings, contrary to the trend observed for sSC neurons. This is consistent with previous findings that responses to opposite motion are potentiated more strongly for sSC neurons than for RGCs^10,41^, suggesting that processing by circuitry intrinsic to the sSC leads to the difference observed here.

## Discussion

We used two-photon fluorescence microscopy with AO to image individual boutons on RGC axons within sSC. Response magnitude and orientation tuning of individual boutons were compared between large uniform gratings and stimuli with edges, opposite motion, and orthogonal orientation, which comprise feature discontinuities within the visual field and hence are salient in a bottom-up manner^6,14–16,42,43^. We found that the majority of RGC boutons responded more strongly to salient stimuli compared to large uniform gratings. In addition, salient stimuli induced dense orientation tuning in RGC boutons, with preferred orientation dictated by the properties of the salient stimulus. These results are similar to our prior observations in sSC neurons^11^ and suggest that saliency encoding in sSC is largely inherited from retinal inputs.

### Orientation selectivity of RGC boutons is highly flexible

Like sSC neurons, RGC boutons showed higher activity at retinotopic locations where salient visual stimuli were present, with the spatial pattern of their activity encoding a map of salience in visual space. By measuring the tuning curves of individual boutons, we found that their activity was not enhanced uniformly for all grating orientations; instead, the increase in activity was greater for specific orientations. In the presence of an edge, most RGC boutons reshaped their orientation selectivity to maximize responses for gratings that were orthogonal to the edge, i.e. the most conspicuous gratings (**Fig. 3**). In addition, boutons along the junction of gratings moving in opposite or orthogonal directions changed their tuning dramatically between the two stimulus types (**Fig. 6**). Overall, our experiments demonstrate that the orientation selectivity of RGC boutons is highly flexible on a massive scale that has not previously been described. Similar findings in sSC neurons indicate that the flexible tuning observed in RGCs is conveyed to their postsynaptic targets^11,13^.

### Both natively and non-natively OS RGC boutons exhibit inducible orientation selectivity and higher activity with salient stimuli

Prior work on RGCs has indicated that a minority are OS in the presence of large drifting gratings or moving bars, with distinct retinal circuitry underlying differences in preferred orientation^33,44–48^. Indeed, when we used large gratings we found that a minor fraction of boutons was OS, consistent with other studies that have examined the orientation tuning of RGC cell bodies in retina^32^ and RGC boutons in SC^21^. We called these boutons “natively tuned” (**Fig. 4**). On the other hand, most boutons were untuned with large gratings but became tuned in the presence of salient stimuli, which we called “non-natively tuned.” Both natively tuned and non-natively tuned boutons had higher responses to salient stimuli and changed their orientation selectivity in similar proportions to oriented edges, oppositely moving gratings, and orthogonally moving gratings.

However, natively tuned boutons tended to become less well tuned when pushed further away from their native preferred orientation, as evidenced by lower gOSI values. This is consistent with the notion that some RGCs are hardwired for specific orientations, especially vertical and horizontal^33^. In contrast, the gOSI of non-natively tuned boutons increased with salient stimuli, such that most of them acquired a preference for orientation. This accords with an earlier study in rat RGCs, in which oppositely moving gratings induced orientation tuning in RGCs that were untuned with uniform gratings^49^. Non-natively tuned boutons may contribute more significantly to saliency encoding since they are more numerous than natively tuned boutons.

### Center-surround RF structure enables saliency encoding

We suspected that the enhanced activity and flexible tuning we observed with salient stimuli arose from the center-surround structure of RGC RFs, in which lateral inhibition from horizontal and amacrine cells suppresses RGC activity to stimuli outside of the RF center^50^. To assess this, we used a Difference-of-Gaussians model to simulate a circularly symmetric RGC RF with an excitatory center and inhibitory surround (**Fig. 5**). Covering a portion of the inhibitory surround with a dark edge yielded higher response from the RF and induced tuning such that responses were higher for gratings oriented orthogonally to the edge. The response was maximal when the edge was near the border of the inhibitory surround and excitatory center of the RF, with most of the excitatory region stimulated by the grating. Responses were also lower for edges of lighter shade, which we confirmed experimentally. This confirms that center-surround RFs that are not intrinsically tuned can nevertheless lead to orientation tuning of RGCs and switch their preferred orientation in a stimulus-dependent manner, facilitating the detection of salient discontinuities in a visual scene. A similar model for SC neurons also demonstrated these phenomena^39^.

Many studies have utilized the center-surround structure of RFs to explore the effects of contextual modulation of RGC activity, in which a stimulus confined to the RF center is surrounded by an annulus with contrasting features. For example, direction selective RGCs are suppressed when moving stimuli extend beyond the center of their RF, but this suppression diminishes if the surrounding stimulus differs in phase^41,51^, spatial or temporal frequency^51^, or direction of motion^41,52,53^. Moreover, it is known that RGCs can alter their responses to oriented stimuli^49^ and movement direction^52–54^ when these properties differ between the center and surround regions of their receptive fields^14,15,42,43^.

In contrast to these previous studies, the saliency encoding we observed does not require precisely confining contrasting stimulus features to the border between center and surround. In our experiments, salient discontinuities in the stimuli likely fall on different RF areas of individual RGCs depending on their precise location, such as would be the case for more naturalistic stimuli. While our model suggests that activity is maximal when an edge falls just outside the RF center, elevated activity occurs at any edge position where part of the inhibitory surround is obscured while most of the excitatory center is exposed to the stimulus (**Fig. 5d**).

Center-surround antagonism has also been implicated in visual functions such as edge detection and figure-ground segmentation^55–58^. Our finding that orientation tuning can be induced in a majority of RGCs and is flexible depending on the properties of visual stimuli suggests that these functions may not be limited to a specialized subset of RGCs, but rather are implemented by a large majority. In addition, flexible orientation selectivity would facilitate the detection of salient stimuli anywhere in the visual field, regardless of regional differences in native orientation preference described recently^21^.

### Potential roles of other contributors to saliency encoding in sSC

Once signals from retina reach the sSC, they undergo further computations via interconnections amongst collicular neurons, which comprise various morphological types, response properties, and projection patterns^59–63^. Circuitry within sSC may further enhance the center-surround properties inherited from retina, and hence amplify saliency encoding^59,64–66^. Some saliency-related signals have been found to differ between different subtypes of retino-recipient sSC neurons^10,13^. Moreover, responses to oppositely moving gratings are much more potentiated for sSC neurons compared to RGCs^10,41^. This is consistent with our observations that RGC boutons tend to respond more for orthogonal gratings than for oppositely moving gratings (**Fig. 6m**), whereas Liang et al.^11^ found the opposite for collicular neurons.

Even though our results suggest that the retina is the main driver of bottom-up saliency in mouse sSC, inputs from visual cortex have also been found to affect the representation of salience within sSC. Removal or silencing of cortical input reduces the response to looming objects in sSC neurons^25^, enhances the “pop-out” effect of orthogonal gratings^26^, and diminishes behavioral responses to a flashing light^27^. Since we removed ipsilateral V1 in our mice, our observations presumably do not depend on cortical feedback; however, this does not exclude effects from sparse contralateral V1 or higher-order cortical inputs, nor the possibility that connections from V1 play a role in other aspects of saliency encoding such as top-down salience, modulation by distractors, or “winner take all” dynamics^6,67–71^. Lastly, the release of neurotransmitters or neuromodulators within sSC, which has receptors for acetylcholine, dopamine, serotonin, and potentially other chemical messengers^72–79^, may also play a role in modulating saliency responses.

### Origin and mechanism of saliency encoding in primate versus rodent

Our work also offers insight into potential differences in saliency encoding across species. Rodents have a prominent SC that receives input from ∼85% of RGCs^8^ and is thought to be considerably more involved in visual processing compared to primate SC^3,55,80–86^. The bottom-up saliency signals we observed in sSC are largely absent from V1^11^, implying that the SC, rather than cortex, plays the most prominent role in behaviors driven by bottom-up saliency in rodents. We further demonstrate here that RGCs in the mouse exhibit many of the same responses as sSC neurons to salient stimuli, suggesting that the sSC derives much of its saliency-related activity from retina.

By contrast, primates have a relatively small SC, which gets input from a measly 10% of RGCs^87^. Saliency signals in primates are thought to originate in V1, then are fed back to SC via corticotectal projections^16–19,67,88^. There is some evidence, however, that saliency-related activity may arise earlier in sSC than cortex^20^, and monkeys with V1 removed still move their eyes toward salient regions of natural scenes^89^. In addition, many primate RGCs have a center-surround RF structure^56,90–92^, which may facilitate the detection of salience through the same mechanisms described here. To our knowledge, stimuli such as the ones used in our study have not been tested in primate retina. Hence the origin of saliency encoding in primate SC, and in particular the involvement of retina, may warrant further study.

Taken together, our results suggest that in mouse, saliency encoding – at least for the stimuli used in this study – originates in the retina. While there has been speculation that retinal inputs establish the salience map in sSC^10,11,13^, this is the first time, to our knowledge, that this conjecture has been experimentally confirmed. Furthermore, since the vast majority of RGCs project to sSC in mouse, and a high proportion of RGC boutons showed heightened activity and flexible orientation selectivity in response to salient stimuli, it follows that saliency encoding is a major function of the mouse retina. At the same time, the contributions of other brain areas – visual cortex, the sSC itself, and neuromodulatory nuclei – is a promising direction for future research.

## Methods

All experimental protocols were done according to the National Institutes of Health guidelines for animal research, and were approved by the Institutional Animal Care and Use Committee at the Janelia Research Campus of Howard Hughes Medical Institute and the University of California, Berkeley.

### Mice

Adult wildtype C57BL/6 mice of both sexes were used for all experiments. Mice were housed in an animal facility with a 12-hour light/dark cycle, ambient temperature 20-26°C, 40-60% humidity.

### Cranial window implantation

Mice were anesthetized with isoflurane (1.5-3% by volume in O_2_) and injected with buprenorphine (intraperitoneal, 0.1 mg per kg body weight) and dexamethasone (subcutaneous, 2.5 mg/kg). When fully anesthetized, mice were placed in a stereotax, lidocaine (3 mg/kg) was injected under the scalp, and eyes were covered with ophthalmic ointment. The skin on top of the skull was shaved and cut away, and the bone surface was cleaned with 2% hydrogen peroxide. To prevent movement of the skull plates during drilling, UV-curing glue (OptiBond Solo Plus, Kerr) was spread in a thin layer on the skull and exposed to UV light for 1 minute. A 3.5-mm diameter craniotomy was drilled over left superior colliculus (SC). The left transverse sinus was tied with two pieces of 8-0 nylon suture thread (AD Surgical). One thread was tied near the lateral edge of the craniotomy, while the other thread was tied near the junction of the transverse and central sinuses. The left transverse sinus was then cut between the two knotted threads. Cortical tissue (including most of V1) was gently aspirated to expose the surface of the SC. Gel foam soaked in cortex buffer was used to protect the exposed brain during surgery (125 mM NaCl, 5 mM KCl, 10 mM glucose, 10 mM HEPES, 2 mM CaCl2, 2 mM MgSO4, 0.125 mg/mL gentamicin, pH 7.4).

Cranial windows were made by gluing a 3 mm diameter glass circle (#1.5 coverslip, Zeiss, laser cut by Potomac Photonics, Inc.) to a 3D-printed plastic barrel (made in-house) with Norland Optical Adhesive (#61 or #68) and cyanoacrylate. Before placing the cranial window onto the mouse, 5 μL of solution containing 2 μm diameter red fluorescent beads (Invitrogen FluoSpheres; diluted in phosphate buffered saline) were pipetted onto the surface of the SC to aid in adaptive optics correction of aberrations. The cranial window was then lowered into the craniotomy and fixed to the skull with cyanoacrylate glue. The 3D-printed barrel had an outer diameter of 3 mm, inner diameter of 2 mm, and was 1.5 mm long. Approximately 1 mm of the barrel extended beneath the skull, and the remainder extended above the surface of the skull. The portion of the barrel above the skull was drilled down a few weeks after surgery, prior to imaging, to be nearly flush with the skull surface.

A titanium or stainless steel head bar was affixed to the exposed skull with Metabond (C&B) and dental acrylic (OrthoJet). Mice were given meloxicam (intraperitoneal, 5-10 mg/kg) on the day of surgery and on the following two post-operative days.

### Intravitreal virus injections

Plasmids and viruses were obtained from the Janelia Research Campus of the Howard Hughes Medical Institute, Addgene, and Vigene Biosciences. Mice were injected intravitreally with recombinant adeno-associated virus to express GCaMP6s either cytosolically (AAV2/2.hSyn.GCaMP6s) or in an axon-targeted manner (AAV2/2.hSyn.GAP43.GCaMP6s)^93^.

Mice were anesthetized with 2% isoflurane, and proparacaine hydrochloride ophthalmic solution (USP 0.5%) was used to anesthetize the right eye. The sclera below the pupil was pierced with a 30-gauge needle, and the tip of a Hamilton syringe was inserted into the puncture to deliver 2 μl of virus. The tip of the syringe was held in the eye for 30 seconds after the injection, then withdrawn quickly. The eye was then covered with ophthalmic ointment and the mouse was allowed to recover.

### Histology

Approximately 4 weeks after intravitreal injection of AAV2/2.hSyn.GAP43.GCaMP6s into the right eye, mice were heavily anesthetized with isoflurane and perfused first with phosphate buffered saline (PBS) for exsanguination, then 4% paraformaldehyde (PFA) for several minutes. The brain and right eye were extracted and incubated in either 4% PFA (brain) or 2% PFA (eye) overnight.

After post-fixation, brains were transferred to a solution of 15% sucrose in PBS overnight, then 30% sucrose in PBS overnight. Brains were then frozen in 30% sucrose on dry ice and sliced in 100 μm coronal sections on a microtome, mounted on microscope slides with VECTASHIELD® HardSet antifade mounting media (Vector Laboratories), and coverslipped.

Eyes were rinsed with PBS and transferred to a petri dish. Retinas were extracted by cutting away the sclera, removing the lens, and separating the retina from the pigmented epithelium. Small cuts were made at a few points along the edge of the retina so it could be laid flat on a microscope slide. The retina was then covered with VECTASHIELD® HardSet antifade mounting media and coverslipped.

Brain slices and retina flatmounts were imaged with a Keyence BZ-X710 fluorescent microscope using either a 4x objective (brain slice) or 10x objective (retina). To create the retina image (**Fig. 1a**), a z-stack was taken at 7 depths, 1 μm apart, and the maximum intensity was calculated across depths to produce the final image.

### Two-photon imaging

Imaging was performed on head-fixed mice > 4 weeks after cranial window implantation. Mice were lightly anesthetized with isoflurane (∼1% in O_2_) during imaging and were lightly restrained with a plastic tube around their body. A goniometer on the head-fixation platform was used to tilt the mouse’s head in order to level the cranial window and minimize optical aberrations.

A femtosecond laser system (InSight Deepsee, Spectra-Physics) tuned to 940 nm was used for fluorescence excitation on a homebuilt two-photon microscope with AO as described previously^29,94^. A Nikon 16x 0.8 N.A. objective was used to deliver excitation light and collect the fluorescence emission. Prior to in vivo imaging, optical aberrations within the microscope’s light path were corrected with AO using a spatial light modulator (SLM; Meadowlark Optics) and 1-μm-diameter red or green beads^94^. When possible, aberrations caused by the mouse’s cranial window were also corrected using the 2-μm-diameter red beads placed under the window during surgery.

In vivo images were acquired at 2.2 Hz with a custom-made program written in LabVIEW. Images typically contained either 300 × 300 pixels, with pixel size being 0.3 or 1.5 µm/pixel for most experiments, or 256 × 256 pixels at 2.3 µm/pixel for retinotopy measurements. In all cases, images were acquired at 256 pixels/ms. The location of the SC surface was estimated by focusing on the 2-μm-diameter red beads under the cranial window. Images of GCaMP6s fluorescence were acquired 40-90 μm below the surface of the SC, with most data acquired at depths of 60-80 μm. Laser power ranged from 20-160 mW measured post-objective, with most data acquired at 50-85 mW. Imaging sessions lasted between 1-6 hours.

### Visual stimulation

Visual stimuli were created in MATLAB (Mathworks) with Psychophysics Toolbox^95^. Illumination from an LED light source (SugarCUBE LED Illuminator; 450-495 nm) and a UV light source (UV 1000-F, ILO Electronic) was combined and delivered via a liquid light guide into a custom-modified DLP projector (Lightspeed DepthQ-WXGA-360; Tan et al. 2015). Stimuli were back-projected onto a Teflon screen (27.5 cm wide, 48 cm tall) placed 13.5 cm from the animal’s right eye, angled 20⁰ relative to the long axis of the animal’s body. The visual display covered 80⁰ of visual space in azimuth and 80⁰ in elevation. The maximal luminance measured at the location of animal eyes was 87 nW/cm^2^ (measured with a power meter calibrated to 400 nm light).

Retinotopy was measured prior to each experiment to find the region of sSC corresponding to the middle of our visual stimulation screen. This enabled us to avoid artifacts induced by the edges of the screen. Retinotopy was measured by flashing light squares on a dark background in a 3 × 3 or 5 × 5 grid in a pseudorandom sequence. The grid covered a total area of 80° visual angle in both azimuth and elevation. Individual squares were either 30° (for the 3 × 3 grid) or 18° (for the 5 × 5 grid) in both dimensions. For each trial, a square was flashed twice (330 ms flash duration, 330 ms between flashes), followed by 4 seconds of darkness before the next trial. Each square position was repeated for 10 trials.

To measure orientation selectivity, sinusoidal gratings moving in one of 12 directions (0° to 330°, in 30° increments) were presented in a pseudorandom sequence. Each direction was repeated for 6 trials. In each trial, a static grating appeared for 4 sec, then drifted for 8 sec. The drifting period was used to measure the response of ROIs (see “Analysis of two-photon imaging data” below). In some experiments the static grating lasted for 5-8 sec, and the drifting period lasted for 7 sec, but this did not affect the magnitude of the response to drifting gratings.

Gratings had 100% contrast and, except for experiments testing different spatial and temporal frequencies, had a spatial frequency of 0.06 cycles per degree and a temporal frequency of 1.54 Hz. Gratings had diameters of 18.5° (small), 52° (mid-size), or 80° (large). Gratings were centered at or near the middle of the screen to avoid artifacts from the screen’s edges.

For experiments testing different spatial frequencies, we used gratings with spatial frequencies of 0.01, 0.02, 0.04, 0.06 (default), 0.08, 0.16, and 0.32 cycles per degree. For experiments testing different temporal frequencies, we used gratings drifting at 0.5, 1, 1.54 (default), 2, 4, 8, and 16 Hz.

For experiments with straight edges, half of the grating was covered by a black uniform rectangle. For experiments with gray edges, the darkness of the uniform rectangle was set to either 30%, 60%, or 100% black. For experiments with oppositely moving or orthogonal gratings, the center of a large (80°) grating was covered by a small (18.5°) grating. The direction of movement of the small grating was rotated either 90° (for orthogonal) or 180° (for oppositely moving) relative to the drifting direction of the large grating. To avoid artifacts from the edge of our stimulus screen, we always placed the edge or small grating at or near the center of the screen and focused on the area of sSC that corresponded retinotopically to that part of the visual field.

### Analysis of two-photon imaging data

ImageJ and custom MATLAB scripts were used to process 2P imaging data^96^. Brain motion was corrected by registering images with an iterative cross-correlation-based rigid registration algorithm^97,98^. ΔF/F_0_ (%) was calculated as (F-F_0_)/F_0_ • 100, where F is the brightness of a pixel or ROI at a given timepoint (frame), and F_0_ is the baseline fluorescence.

Retinotopy mapping: Images were Gaussian filtered (filter size = 20 pixels, sigma = 20 pixels). Brightness F of each pixel was extracted and averaged across trials (10 trials for each retinotopy grid position). The trial-averaged F of each pixel was then tested with ANOVA across stimulus locations in the retinotopy grid (cutoff p = 0.1). Pixels that passed the statistical test were assigned to a location on the retinotopy map according to the stimulus that elicited the highest response from that pixel.

Pixel-based analysis of maximum response: Baseline fluorescence F_0_ was calculated as the mode of fluorescence values across all frames of a given session (12 grating directions, 6 repeats per direction) and used to calculate ΔF/F_0_. For each grating direction, ΔF/F_0_ was averaged across frames during stimulus movement in each trial (excluding the first five frames after movement onset to allow fluorescence to reach its maximum value). Frame-averaged fluorescence was then averaged across trials for each grating direction. The highest value of ΔF/F_0_ across the 12 stimulus angles was defined as the maximum response.

Pixel-based analysis of orientation selectivity: Orientation selectivity was determined for each pixel in the image. Images were Gaussian filtered (filter size = 3 pixels, sigma = 1 pixel). Baseline fluorescence F_0_ was calculated as the mode of fluorescence values across all frames of a given session (12 grating directions, 6 repeats per direction). Negative fluorescence values were assumed to be noise, and values were set to 2.22e-16. To determine the response of each pixel for a particular grating direction, fluorescence was first averaged during the period of grating movement, omitting the first 5 frames after grating movement onset to allow the calcium signal to reach its maximum level. Fluorescence was then averaged across trials for a given direction.

Pixels that passed a one-way ANOVA test (with the response to at least one stimulus angle being significantly different from the other angles, p < 0.05) were designated as orientation selective. To determine the preferred orientation of a given pixel, responses were fit to a double Gaussian equation:

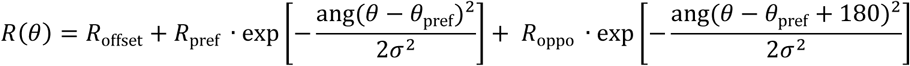

*R*(*θ*) is the response at stimulus angle *θ*, *R*_offset_ is a constant, *R*_pref_ is the response at *θ*_pref_, and *R*_oppo_ is the response at *θ*_pref_ – 180°. The function ang wraps values for stimulus angle from 0° to 180°, ang(*θ*) = min(*θ*, *θ* – 360°, x + 360°).

ROI analysis of orientation selectivity and direction selectivity: For high resolution images, ROIs were drawn either manually in ImageJ or with Suite2p^31^. ROIs drawn in Suite2p were compared against either hand-drawn ROIs or pixel-based analysis to verify accuracy, and were hand-curated to eliminate spurious ROIs. Fluorescence was averaged for all pixels within a given ROI, for each frame. Baseline F_0_ was the average fluorescence during the last two frames prior to the onset of grating movement and was determined on a per-trial basis. ROIs were classified as being visually responsive if maximum ΔF/F_0_ > 10% for at least one visual stimulus tested.

To calculate global orientation index (gOSI) for a given ROI, we used the following equation:

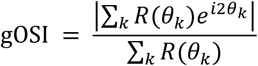

where *R*(*θ*_*k*_) is the response towards sinusoidal gratings moving along angle *θ*_*k*_, with *θ*_*k*_ being one of 12 directions (0° to 330°, in 30° increments).

To calculate global direction index (gDSI) for a given ROI, we used the following equation:

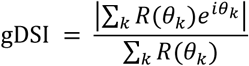

To determine whether a given ROI was OS, we used a shuffling algorithm^32^. The relation between stimulus angle and response was randomized, gOSI was calculated, and this was repeated 1000 times to generate a null distribution of gOSI values. The actual gOSI value of the ROI was then compared against this distribution. If the actual gOSI value fell outside the 95^th^ percentile of the null distribution (p < 0.05), the ROI was considered to be OS. This method avoids assigning OS status to ROIs that may have a high gOSI, but whose responses are too variable to be reliably orientation tuned. To determine the preferred orientation of an ROI, responses were fit to the same double Gaussian equation shown above for pixel analysis.

To determine whether a given ROI was DS, we performed the same shuffling algorithm with gDSI values. Non-DS ROIs that failed our statistical test were not included in the DS population even if they had a high gDSI. Preferred direction was calculated as the vector sum of responses across the 12 stimulus directions.

ROI analysis of maximum response: ROIs were included for analysis if they were responsive to at least one of the stimuli presented (maximum ΔF/F_0_ > 10%). The maximum value of response R(*θ*) across the 12 stimulus angles was defined as the maximum response.

ROI analysis of gOSI: For cumulative distributions of gOSI, the dataset for each stimulus included ROIs that were responsive for that stimulus. For scatterplots, ROIs were included if they were responsive to both stimuli being compared.

ROI analysis of preferred orientation: For distributions of preferred orientation, the dataset for each stimulus included ROIs that were OS for that stimulus. For distributions of change in preferred orientation, ROIs were included if they were OS to both stimuli being compared.

To calculate the percentage of natively tuned ROIs that prefer vertical or horizontal gratings under different edge conditions (see Results for **Fig. 4**), we considered each ROI to prefer a particular orientation if its fitted angle fell within a range ± 20⁰ of the designated orientation. For example, an ROI was considered to prefer vertical or near-vertical gratings if its fitted angle fell within the range 70⁰ - 110⁰, and was considered to prefer horizontal or near-horizontal gratings if its fitted angle fell within the range 0⁰ - 20⁰ or 160⁰ - 180⁰.

Statistical tests: For testing significance of differences between cumulative distributions and of gOSI, gDSI, and max dF/F_0_, we used the two-sample Kolmogorov-Smirnov test. For testing significance of differences in percentage of OS boutons between different stimulus conditions, we used the Wilcoxon signed-rank test. For testing whether the change in gOSI or change in max dF/F_0_ was significantly different from zero, we used the one-sided Wilcoxon sign-rank test (**Supp. Fig 6** and **Supp. Fig. 9**). For testing significance of differences between paired datasets in these figures (e.g. vertical versus horizontal edge), we used the Wilcoxon rank-sum test. All p values were reported in **Supplementary Tables**.

### Model of RGC receptive fields

We used a Difference-Of-Gaussians (DOG) model to simulate the center-surround RFs of retinal ganglion cells. We assumed a circularly symmetric RF with an excitatory center and a concentric inhibitory surround^50^. We calculated excitatory and inhibitory components as two-dimensional Gaussian functions:

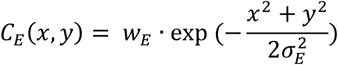

and

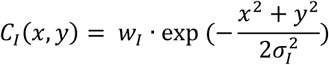

where *w*_*E*_ is the weight and *σ*_*E*_ is the RMS width for *C*_*E*_(*x*, *y*), *w*_*I*_ is the weight and *σ*_*I*_ is the RMS width for *C*_*I*_(*x*, *y*), and *x* and *y* are spatial coordinates. The receptive field *C*(*x*, *y*) was calculated as the difference of the two Gaussian functions:

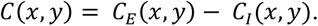

The total response of the RF was calculated as:

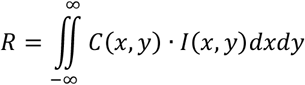

where *I*(*x*, *y*) describes the illumination intensity distribution of a visual stimulus. We calculated the illumination from a sinusoidal grating as follows:

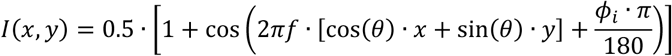

where *f* is the spatial frequency of the grating, *θ* is the orientation of the grating, and *Φ*_*i*_ is the initial phase of the grating. Spatial frequency was set to 0.06 cycles per pixel, matching our experimental parameters. We calculated responses for 12 grating orientations (0° to 330°, in 30° increments). Grating phase was advanced in increments of π/180 in order to simulate a moving grating, and the responses were averaged for phases 0 – 2π. Edges were simulated as areas with *I*(*x*, *y*) = 0 (for black edge) or a nonzero constant value (for gray edges).

We chose *w*_*E*_ = 1.8, *w*_*I*_ = 1, and *σ*_*I*_ = 30 µm in accordance with previously reported experimental values^36–38^. To determine *σ*_*E*_, we set the condition that the net response of the receptive field be 0 with uniform illumination (i.e., *I*(*x*, *y*) = 1), with

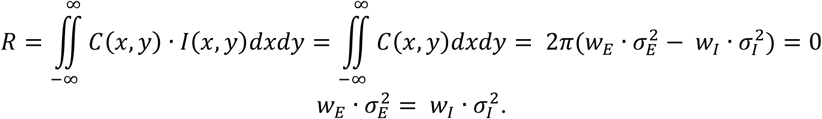

Excitatory weight was therefore calculated as

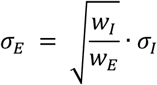

With the above values for *w*_*E*_, *w*_*I*_, and *σ*_*I*_, *σ*_*E*_ was determined to be 0.75*σ*_*I*_ or 22 µm. Analyses in **Fig. 6** were then performed using these parameters.

We also tested the model’s performance with different ratios of *w*_*E*_/*w*_*I*_ and *σ*_*E*_/*σ*_*I*_. Values within the range 1.6 – 2.4 for *w*_*E*_/*w*_*I*_ and 0.7 – 0.8 for *σ*_*E*_/*σ*_*I*_ resulted in similar behavior of the model, with tuning induced by an edge and a preference for gratings orthogonal to the orientation of the edge.

## Acknowledgements

We thank Peijia Yu for contributions to code for Supp. Fig. 1c,d; Xinlu Ding for assistance with imaging of brain slices and retina flatmounts, Wei Chen for technical assistance with the 2P system. This work was supported by Weill Neurohub and National Institutes of Health (U01 NS118300, U01 NS120820).

## Author contributions

N.J. supervised the project. K.B. and N.J. designed experiments. K.B. performed all experiments, wrote custom analysis code, and performed all data analysis. Q.X., Y.L., and R.L. contributed code and preliminary analysis of the Difference-Of-Gaussians model. R.L. and S.H. contributed additional code for data analysis. S.N. contributed manual ROI segmentation. KB and N.J. wrote the paper.

## Supplementary Figures

**Supplementary Figure 1.**
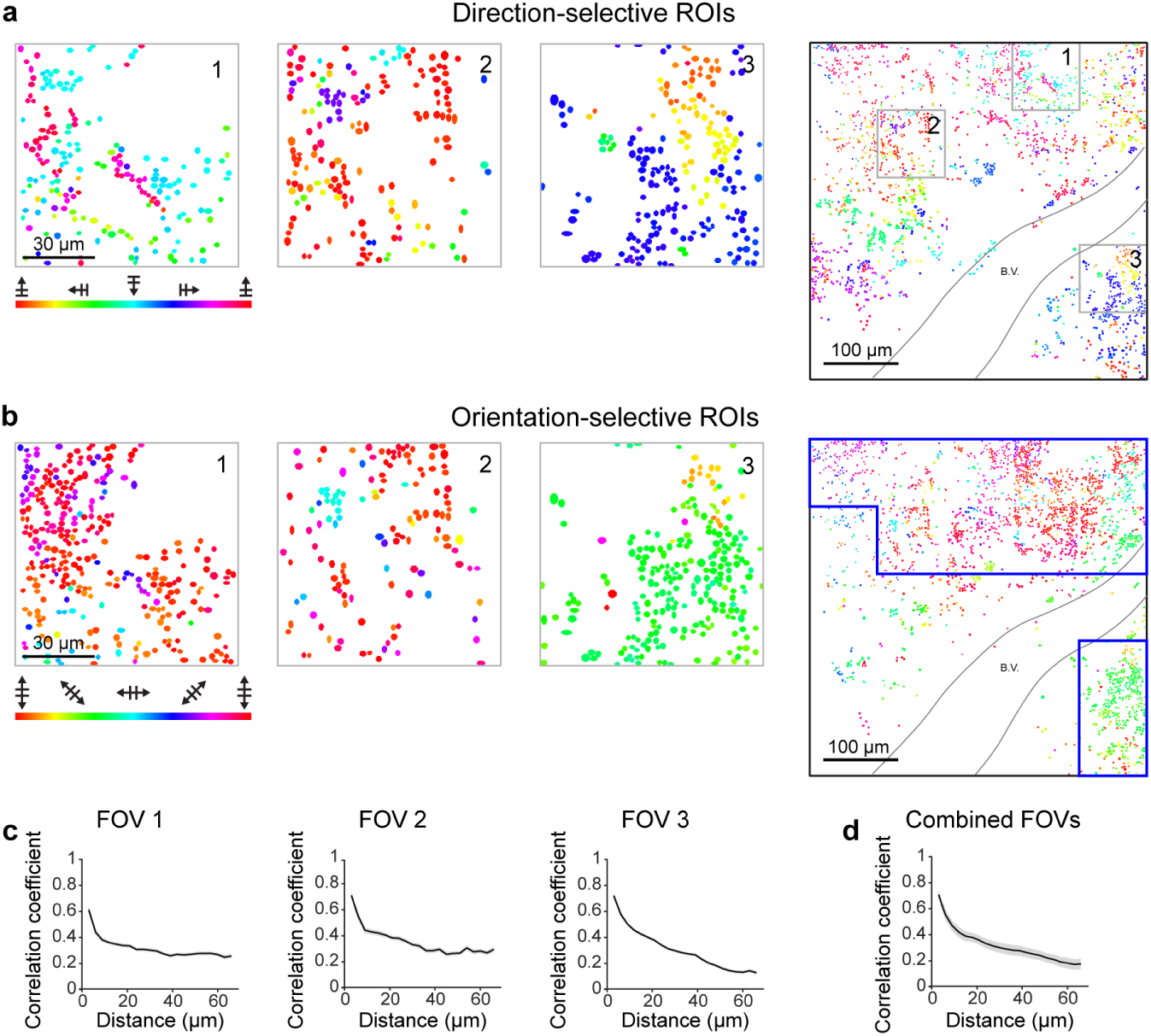
RGC boutons form localized clusters with similar orientation and direction preference. (**a**) Enlarged views of 3 FOVs labeled in the composite of 25 FOVs on the right, with DS boutons color-coded by preferred direction. (**b**) Same as **a**, with OS boutons color-coded by preferred orientation. (**c**) Correlation coefficient between ΔF/F_0_ traces each pair of boutons versus distance between the two boutons within the indicated FOV. Shaded areas: ± S.E.M. (**d**) Same as **c**, for the population of boutons in FOVs with dense tuning outlined in the blue boxes in **b** (11 out of 25 FOVs).

**Supplementary Figure 2.**
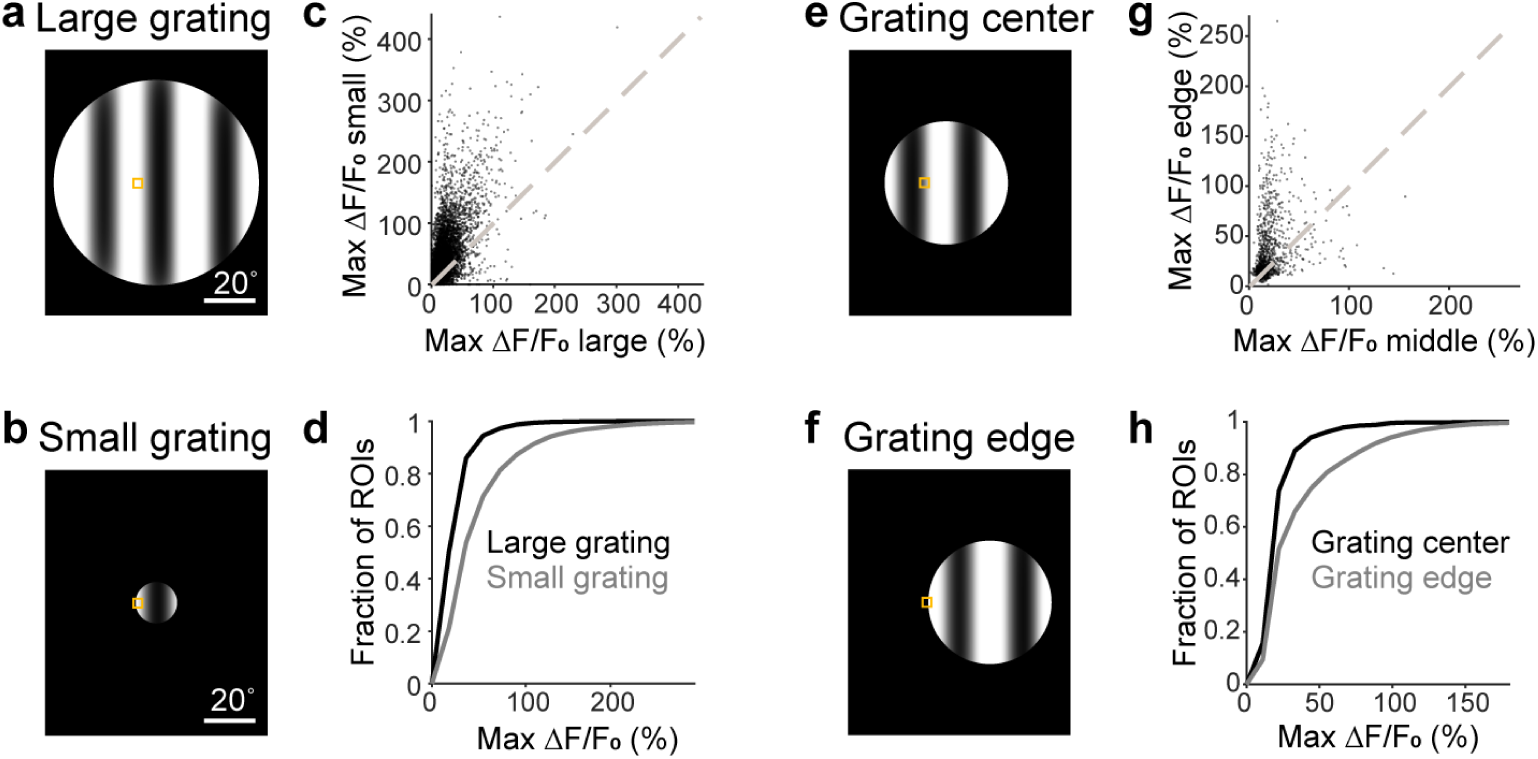
Grating edge enhances calcium activity of RGC boutons. (**a**) Large-aperture drifting grating. Orange box: retinotopic area of FOVs imaged at high resolution. (**b**) Small-aperture drifting grating. (**c**) Scatter plot of max ΔF/F_0_ of 6943 visually responsive boutons from 13 FOV of 8 mice for small versus large gratings. Each datapoint represents one bouton. 81% of boutons above unity line. (**d**) Cumulative distributions of max ΔF/F_0_ for large (black; median: 18.1%) or small (gray; median: 33.4%) gratings, for boutons that are responsive to either grating. *p* value: 0. (**e**) FOVs mapped to the center of drifting gratings. (**f**) FOVs mapped to the edge of drifting gratings. (**g**) Scatter plot of max ΔF/F_0_ of 1125 visually responsive boutons from 2 FOVs of 2 mice for grating center vs. edge. 67% of boutons above unity line. (**h**) Cumulative distributions of max ΔF/F_0_ for grating center (black; median: 16.3%) and edge (gray; median: 21.6%), for boutons that are responsive to both gratings. Medians are significantly different; *p* value: 7.28e-29.

**Supplementary Figure 3.**
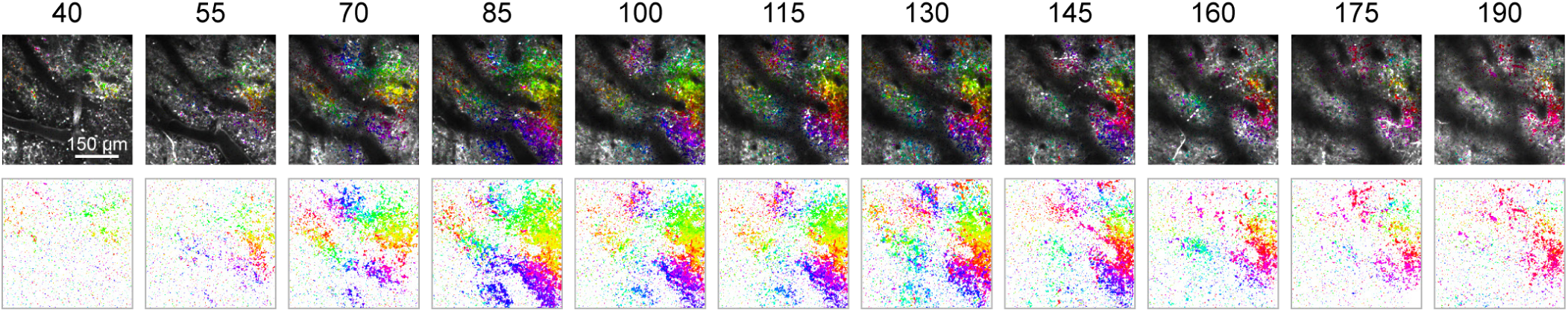
RGC boutons respond with dense orientation tuning to grating edge across depth. 2P images with superimposed orientation selectivity tuning maps (top) and orientation selectivity tuning maps alone (bottom) of FOVs at depths from 40 to 190 μm below the surface of the sSC from one mouse, as indicated by the numbers above each panel. Grating size: 18.5° visual angle. Similar results were obtained in 4 additional mice.

**Supplementary Figure 4.**
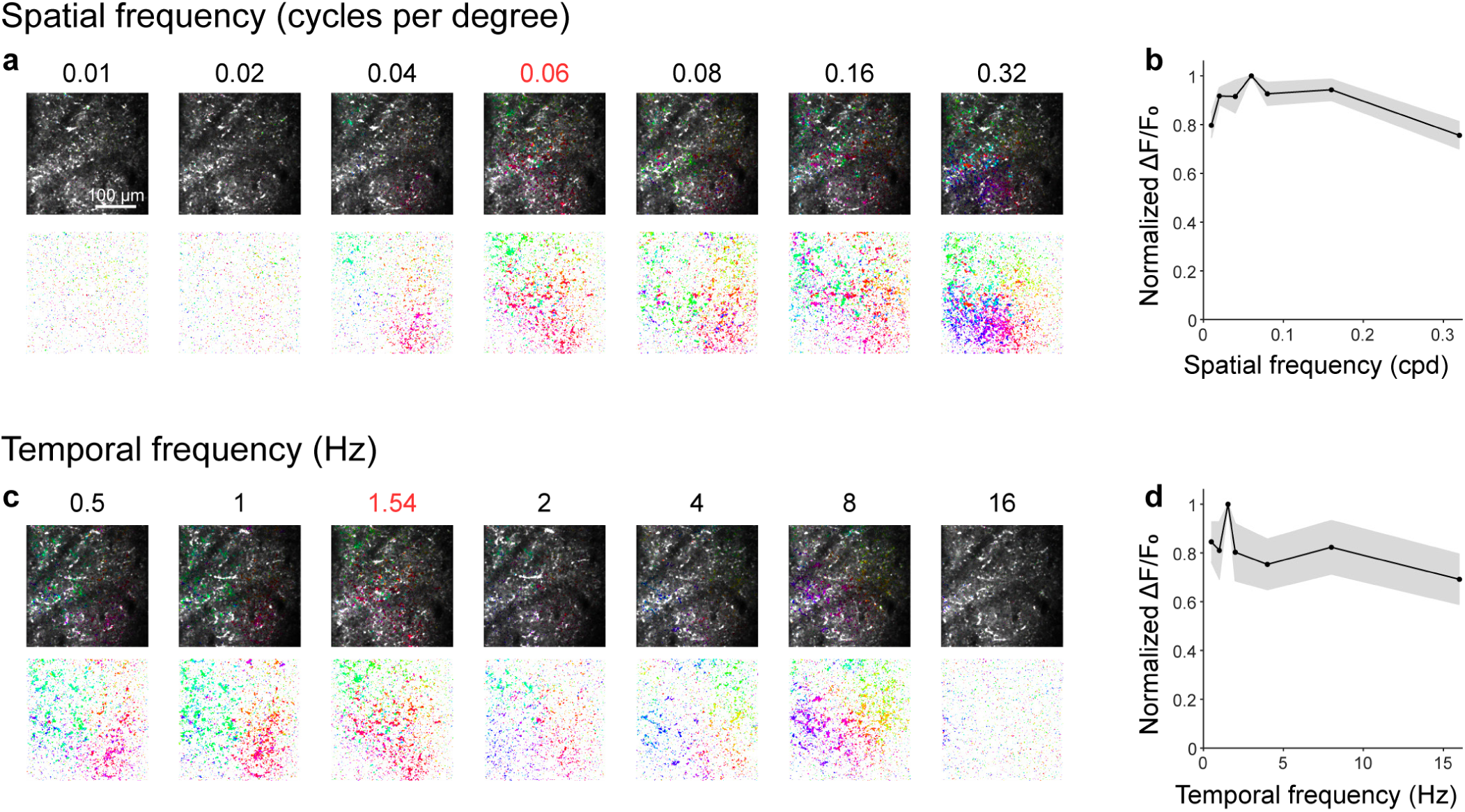
RGC boutons show orientation tuning to grating edge across various spatial and temporal frequencies. (**a**) 2P images with superimposed orientation selectivity maps (top) and orientation selectivity maps alone (bottom) imaged from a FOV while varying the spatial frequency of the grating. Default spatial frequency used in all other experiments is indicated in red. Grating size: 18.5° visual angle. (**b**) Average max ΔF/F_0_ at each spatial frequency for 9 FOVs in 8 mice. Max ΔF/F_0_ was normalized to the default value (0.06 cpd) for each FOV prior to averaging. (**c**) Same as **a**, varying the temporal frequency of the grating. (**d**) Same as **b**, for temporal frequency. Data are from 7 FOVs in 6 mice.

**Supplementary Figure 5.**
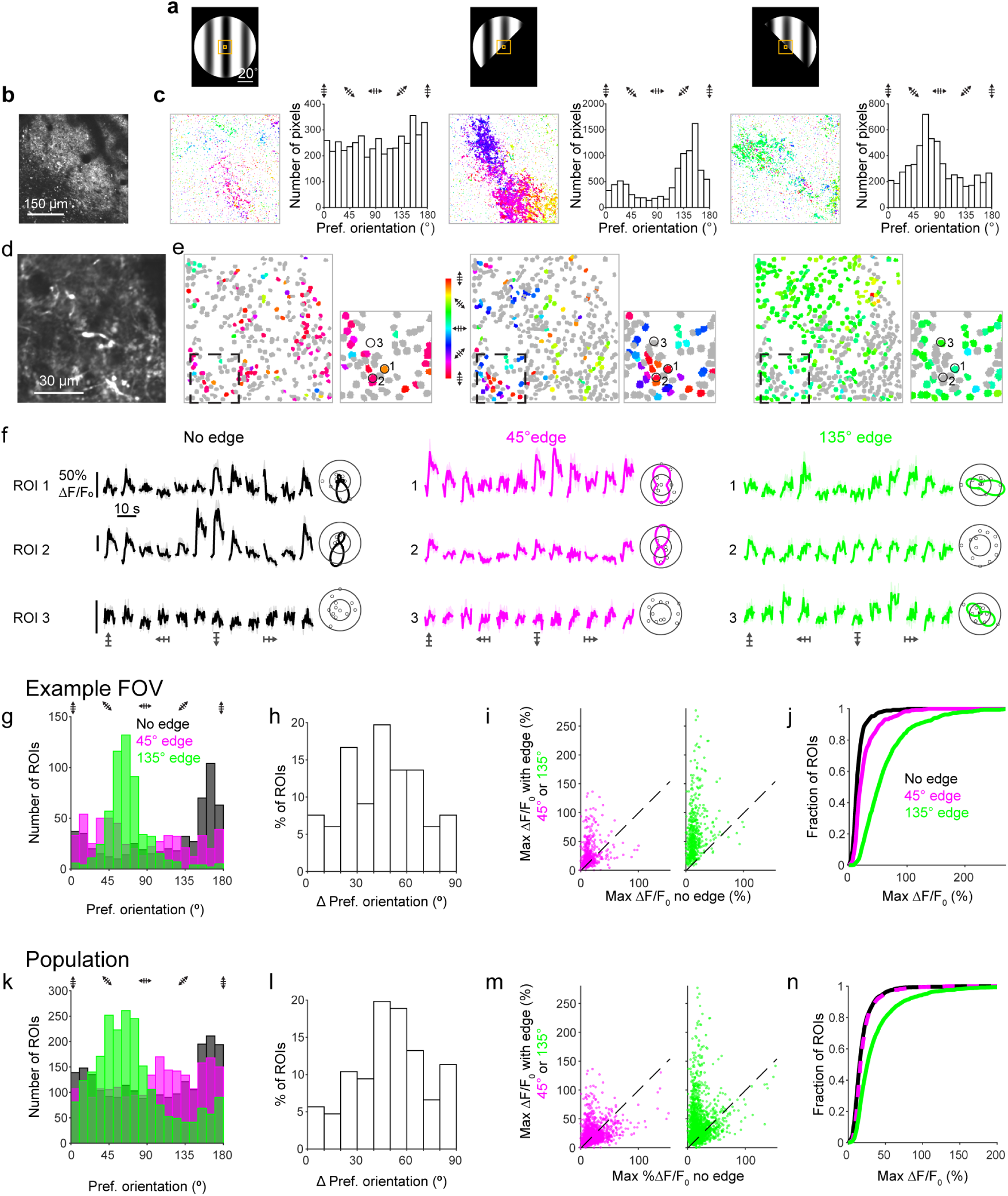
RGC boutons change their tuning to prefer orientations that are perpendicular to oblique edges. (**a**) Stimuli: large drifting grating (80° visual angle) with either no edge (left) or with a straight 45° (middle) or 135° (right) edge covering half the grating. (**b**) 2P image of RGC boutons in a 450 μm × 450 μm FOV in sSC. Pixel size: 1.5 μm. (**c**) OS pixels color-coded by preferred orientation (left) and preferred orientation distributions (right) for pixels in **b**, for each stimulus in **a**. (**d**) Image of RGC boutons in an example 90 μm × 90 μm FOV in sSC. Pixel size: 0.3 μm. (**e**) RGC boutons from **d**, color-coded based on orientation selectivity in response to stimuli in **a**. Gray: visually responsive but not OS boutons; colored: OS boutons. Insets: Enlarged views of boutons in dashed boxes. (**f**) ΔF/F_0_ traces and polar plots for the three circled ROIs in **e**, for each stimulus in **a**. Shaded areas: ± S.E.M. (**g**) Preferred orientation distributions for boutons in **d** that are OS with no edge (black), 45° edge (magenta), and 135° edge (green). (**h**) Change in preferred orientation between 45°and 135°edges, for boutons in **d** that are OS with both edges. (**i**) Scatter plots of max ΔF/F_0_ for visually responsive boutons in **d** with 45° (left) or 135° (right) edge vs. no edge. 71% and 96% of boutons above unity line for 45° and 135° edges, respectively. (**j**) Cumulative distributions of max ΔF/F_0_ for visually responsive boutons in **d** for each stimulus. Medians for no edge, 45° edge, 135° edge: 12.8%, 19.0%, 50.5%, respectively. *p* values: 2.10e-19 for no edge vs. 45° edge, 2.14e-130 for no edge vs. 135° edge, 10.0e-68 for 45° vs. 135° edge. (**k-n**) Same as **g-j**, for a population of 2192 boutons in 4 FOVs from 3 mice. **m**: 54% and 73% of boutons above unity line for 45° and 135° edges, respectively. **n**: Medians for no edge, 45° edge, 135° edge: 15.2%, 16.4%, 25.7%, respectively. *p* values: 1.45e-9 for no edge vs. 45° edge, 3.26e-106 for no edge vs. 135° edge, 9.28e-85 for 45° vs. 135° edges.

**Supplementary Figure 6.**
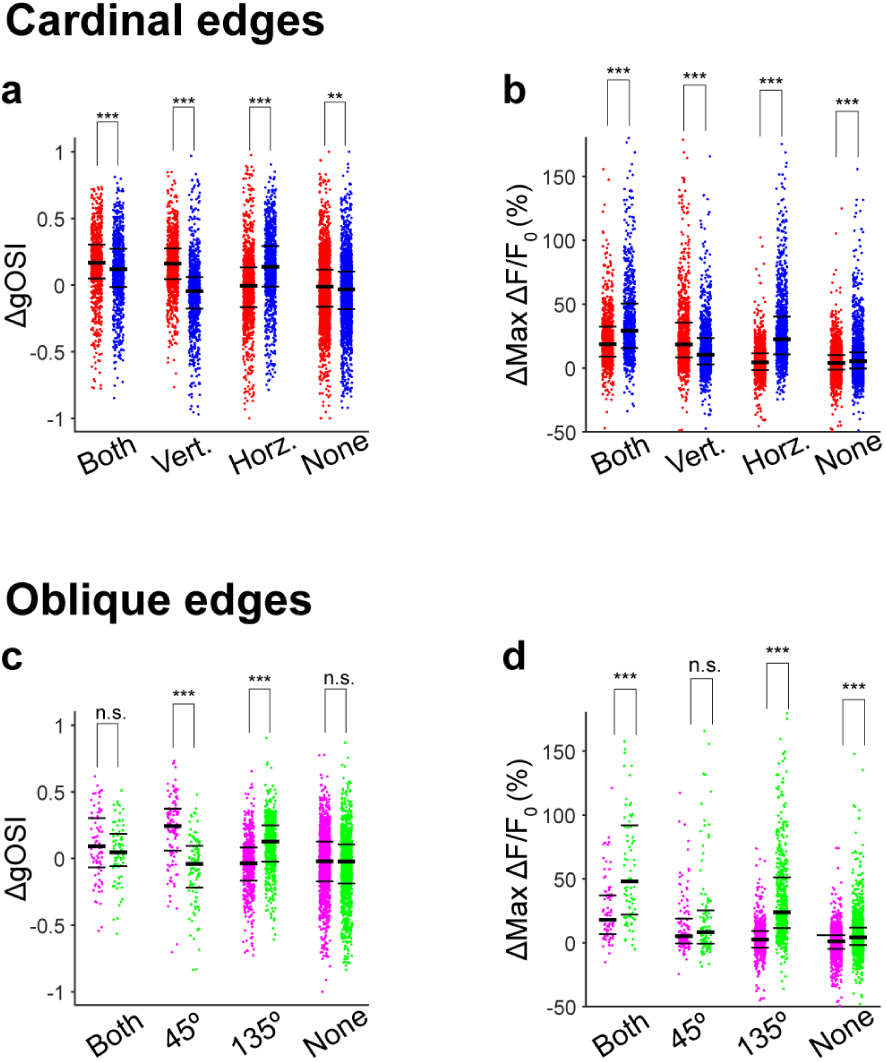
Changes in gOSI and max ΔF/F_0_ for non-natively tuned boutons. Analysis for the same datasets as Fig. 3 and **Supp.Fig. 5**. (**a**) Change in gOSI between no-edge and edge conditions, for non-natively tuned boutons that are tuned for (pairs, left to right) both edges, vertical edge only, horizontal edge only, or neither edge. Each of the four pairs shows ΔgOSI for vertical edge versus no edge (red dots) and horizontal edge versus no edge (blue dots), for the same subset of boutons. See **Supp. Table 7** for all medians and *p* values. (**b**) Change in max ΔF/F_0_ between no-edge and edge conditions for non-natively tuned boutons, grouped as in (**a**). (**c-d**) Same as **a-b**, for oblique edges (magenta: 45°, green: 135°).

**Supplementary Figure 7.**
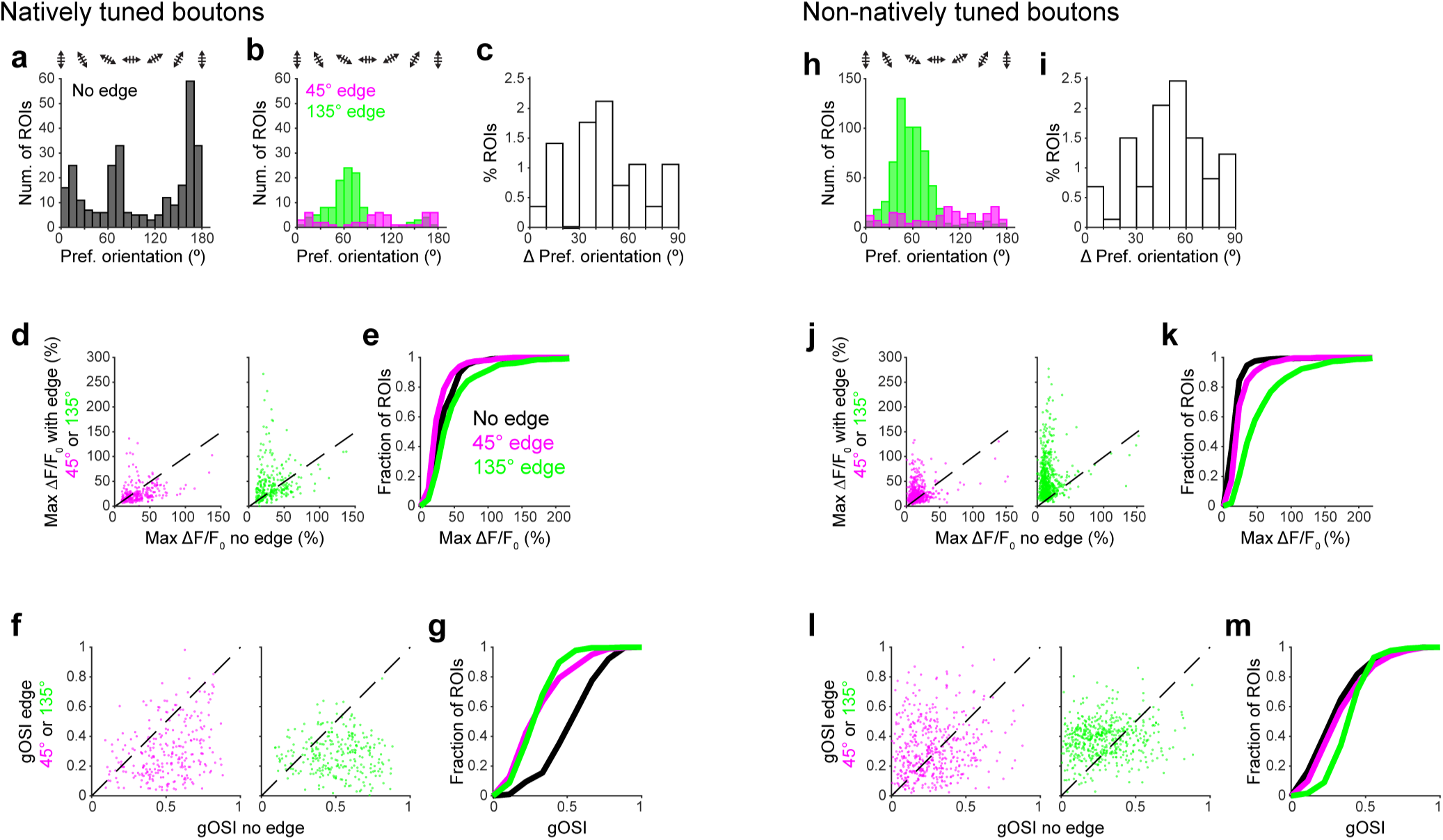
Oblique edges modify the orientation tuning properties of both natively tuned and non-natively tuned RGC boutons. Data are from the same mice as Supplemental Fig. 5k-n. (**a**) Preferred orientation distribution for natively tuned boutons with large grating (no edge). (**b**) Same as **a**, for 45° (magenta) and 135° (green) edges. (**c**) Change in preferred orientation between 45°and 135°edges, for natively tuned boutons that are OS with both edges. (**d**) Scatter plots of max ΔF/F_0_ for natively tuned boutons with 45° (left) or 135° (right) edge versus no edge. 28% and 54% of boutons above unity line for 45° and 135°, respectively. (**e**) Cumulative distributions of max ΔF/F_0_ for natively tuned boutons, for each stimulus. Medians for no edge, 45° edge, and 135° edge: 30.0%, 20.7%, 34.9%, respectively. *p* values: 2.49e-8 for no edge vs. 45° edge, 4.88e-4 for no edge vs. 135° edge, 4.27e-20 for 45° vs. 135° edge. (**f**) Scatter plots of gOSI for natively tuned boutons with 45° (left) or 135° (right) edge vs. no edge. 17% boutons above unity line for both 45° and 135° edges. (**g**) Cumulative distributions of gOSI for natively tuned boutons. Medians for no edge, 45° edge, and 135° edge: 0.52, 0.27, 0.28, respectively. *p* values: 2.69e-30 for no edge vs. 45° edge, 1.01e-39 for no edge vs. 135° edge, 0.0255 for 45° vs. 135° edge. (**h-m**) Same as **b-g**, for non-natively tuned boutons. **j**: 66% and 90% of boutons above unity line for 45° and 135°, respectively. **k**: Medians for no edge, 45° edge, and 135° edge: 13.3%, 18.1%, 40.4%, respectively. *p* values: 1.69e-19 for no edge vs. 45° edge, 3.06e-136 for no edge vs. 135° edge, 1.03e-76 for 45° vs. 135° edge. **l**: 55% and 70% of boutons above unity line for 45° and 135°, respectively. **m**: Medians for no edge, 45° edge, and 135° edge: 0.26, 0.29, 0.39, respectively. *p* values: 0.0298 for no edge vs. 45° edge, 1.26e-34 for no edge vs. 135° edge, 9.21e-26 for 45° vs. 135° edge.

**Supplementary Figure 8.**
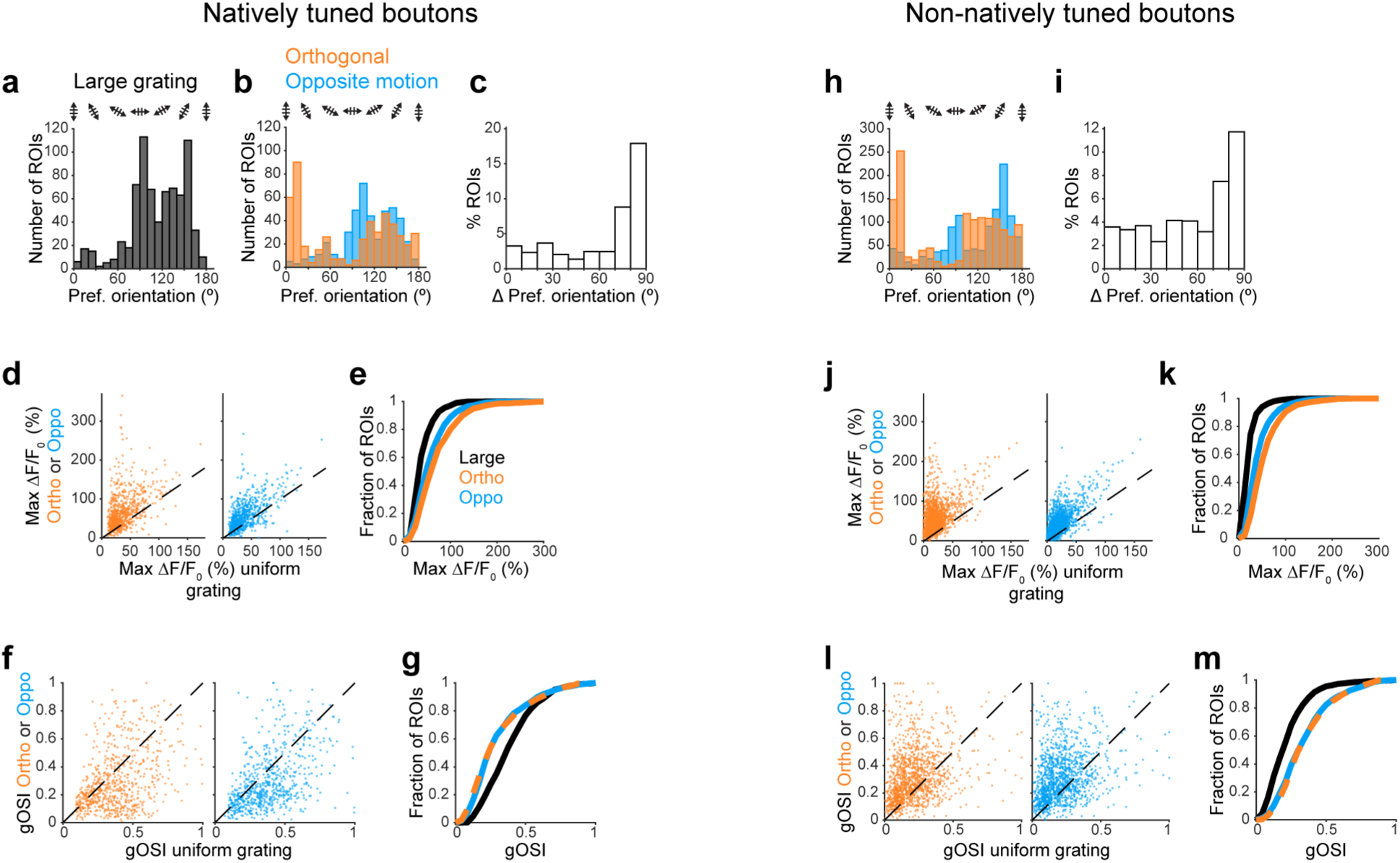
Border between concentric gratings modifies the orientation tuning properties of natively tuned and non-natively tuned RGC boutons. (**a**) Preferred orientation distribution for natively tuned boutons with large grating. (**b**) Same as **a**, for orthogonal gratings (OG, orange) and oppositely moving gratings (OMG, blue). (**c**) Change in preferred orientation between OG and OMG, for natively tuned boutons that are OS with both stimuli. (**d**) Scatter plots of max ΔF/F_0_ for natively tuned boutons with OG (left) or OMG (right) versus no uniform gratings. 78% and 82% of boutons above unity line for OG and OMG, respectively. (**e**) Cumulative distributions of max ΔF/F_0_ for natively tuned boutons, for each stimulus. Medians for large grating, OG, OMG: 32.2%, 57.5%, 47.8%, respectively. *p* values: 1.31e-42 for large vs. ortho. gratings, 4.27e-23 for large vs. oppo. gratings, 1.80e-5 for ortho. vs. oppo. gratings. (**f**) Scatter plots of gOSI for natively tuned boutons with OG (left) or OMG (right) vs. large gratings. 34% and 30% of boutons above unity line for OG and OMG, respectively. (**g**) Cumulative distributions of gOSI for natively tuned boutons. Medians for large grating, OG, OMG: 0.36, 0.23, 0.23, respectively. *p* values: 1.63e-24 for large vs. ortho. gratings, 7.71e-25 for large vs. oppo. gratings, 0.1454 for ortho. vs. oppo. gratings. (**h-m**) Same as **b-g**, for non-natively tuned boutons. **j**: 95% and 92% of boutons above unity line for OG and OMG, respectively. **k**: Medians for large grating, OG, OMG: 16.2%, 45.8%, 33.0%, respectively. *p* values: 7.48e-271 for large vs. ortho. gratings, 5.55e-143 for large vs. oppo. gratings, 2.16e-34 for ortho. vs. oppo. gratings. **l**: 73% and 73% of boutons above unity line for OG and OMG, respectively. **m**: Medians for large grating, OG, OMG: 0.19, 0.30, 0.30, respectively. *p* values: 2.40e-68 for large vs. ortho. gratings, 8.77e-56 for large vs. oppo. gratings, 0.0911 for ortho. vs. oppo. gratings.

**Supplementary Figure 9.**
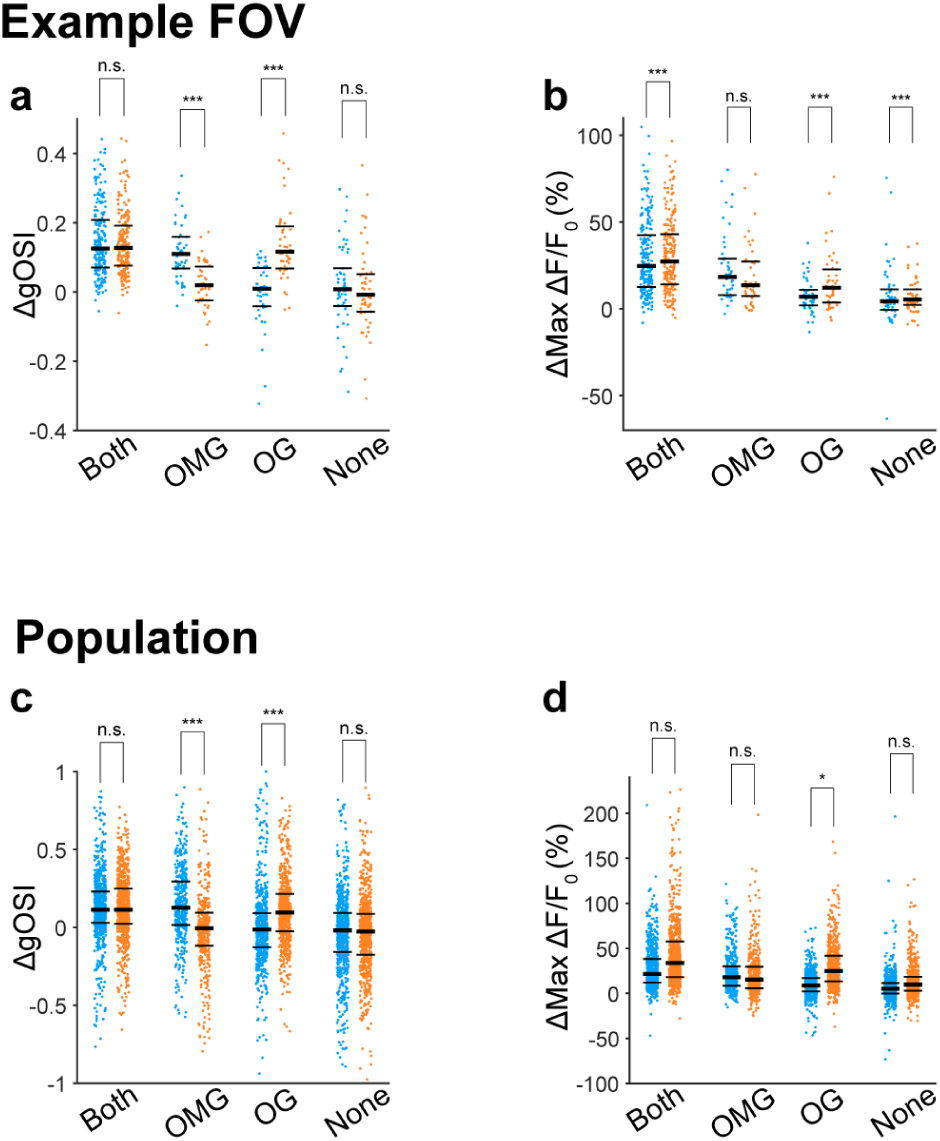
Changes in gOSI and max ΔF/F_0_ for non-natively tuned boutons in response to oppositely moving and orthogonal gratings. (**a-b**) Analysis for the FOV shown in Fig. 7d: (**a**) Change in gOSI between a uniform grating and OMG or OG, for non-natively tuned boutons that are tuned for (pairs, from left to right) both OMG and OG, OMG only, OG only, or neither of the concentric gratings. Each of the four pairs shows ΔgOSI for OMG versus large grating (blue dots) and OG versus large grating (orange dots), for the same subset of boutons. See **Supp. Table 12** for all medians and *p* values. (**b**) Change in max ΔF/F_0_ between large grating and OMG or OG for non-natively tuned boutons, grouped as in (**a**). (**c-d**) Same as **a-b**, for a population of 1108 boutons, 6 FOVs, 4 mice.

## Supplementary Tables

**Supplementary Table 1:** gOSI and gDSI distributions for Fig. 1.

| Panel h: gDSI distributions |  |  |  |  |
| --- | --- | --- | --- | --- |
| Bouton type | # boutons | median gDSI | significantly different | p value |
| Direction selective | 2268 | 0.43 | yes | 8.63e-154 |
| Responsive non-DS | 4742 | 0.19 |  |  |
| Panel k: gOSI distributions |  |  |  |  |
| Bouton type | # boutons | median gOSI | significantly different | p value |
| Orientation selective | 2374 | 0.33 | yes | 9.89e-80 |
| Responsive non-OS | 4636 | 0.22 |  |  |

**Supplementary Table 2:** Percent OS ROIs and max ΔF/F_0_ distributions for Fig. 2.

| Panel e,f: Percent OS ROIs, example FOV |  |  |  |  |
| --- | --- | --- | --- | --- |
| Stimulus | # responsive boutons | # OS boutons | % OS |  |
| Large grating | 436 | 128 | 29 |  |
| Small grating | 483 | 213 | 44 |  |
| Panel h: Max ΔF/F<sub>0</sub> distributions, example FOV |  |  |  |  |
| Stimulus | # boutons (responsive to at least one stimulus) | median max ΔF/F<sub>0</sub> | significantly different from | p value |
| Large grating | 491 | 19.2% | Small grating: yes | 2.46e-59 |
| Small grating | 491 | 44.5% |  |  |
| Panel k,l: Percent OS ROIs, example FOV |  |  |  |  |
| Stimulus | # responsive boutons | # OS boutons | % OS |  |
| Middle of grating | 499 | 81 | 16 |  |
| Edge of grating | 510 | 240 | 47 |  |
| Panel n: Max ΔF/F<sub>0</sub> distributions, example FOV |  |  |  |  |
| Stimulus | # boutons (responsive to at least one stimulus) | median max ΔF/F<sub>0</sub> | significantly different from | p value |
| Middle of grating | 522 | 19.1% | Edge of grating: yes | 1.60e-21 |
| Edge of grating | 522 | 30.4% |  |  |

**Supplementary Table 3:** Percent OS ROIs and max ΔF/F_0_ distributions for Supp. Fig. 2.

| Panel d: Max $\Delta F/F_0$ distributions, population | | | | |
| --- | --- | --- | --- | --- |
| Stimulus | # boutons<br>(responsive to at<br>least one stimulus) | median max $\Delta F/F_0$ | significantly different<br>from | p value |
| Large grating | 6943 | 18.1% | Small grating: yes | 0 |
| Small grating | 6943 | 33.4% |  |  |
| Panel h: Max $\Delta F/F_0$ distributions, population | | | | |
| Stimulus | # boutons<br>(responsive to at<br>least one stimulus) | median max $\Delta F/F_0$ | significantly different<br>from | p value |
| Middle of grating | 1125 | 16.3% | Edge of grating: yes | 7.28e-29 |
| Edge of grating | 1125 | 21.6% |  |  |

**Supplementary Table 4:**
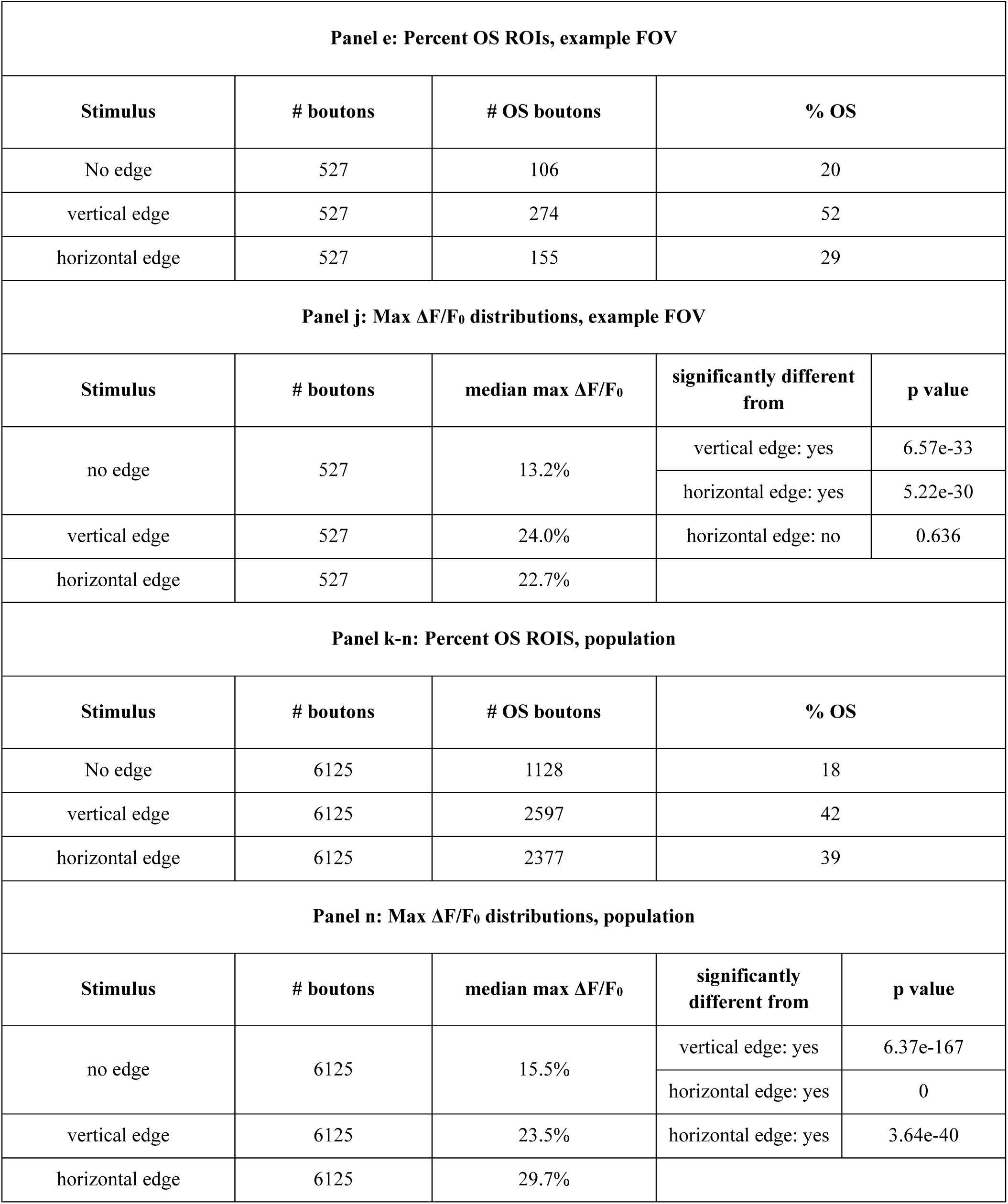
Percent OS ROIs and max ΔF/F_0_ distributions for Fig. 3.

**Supplementary Table 5:**
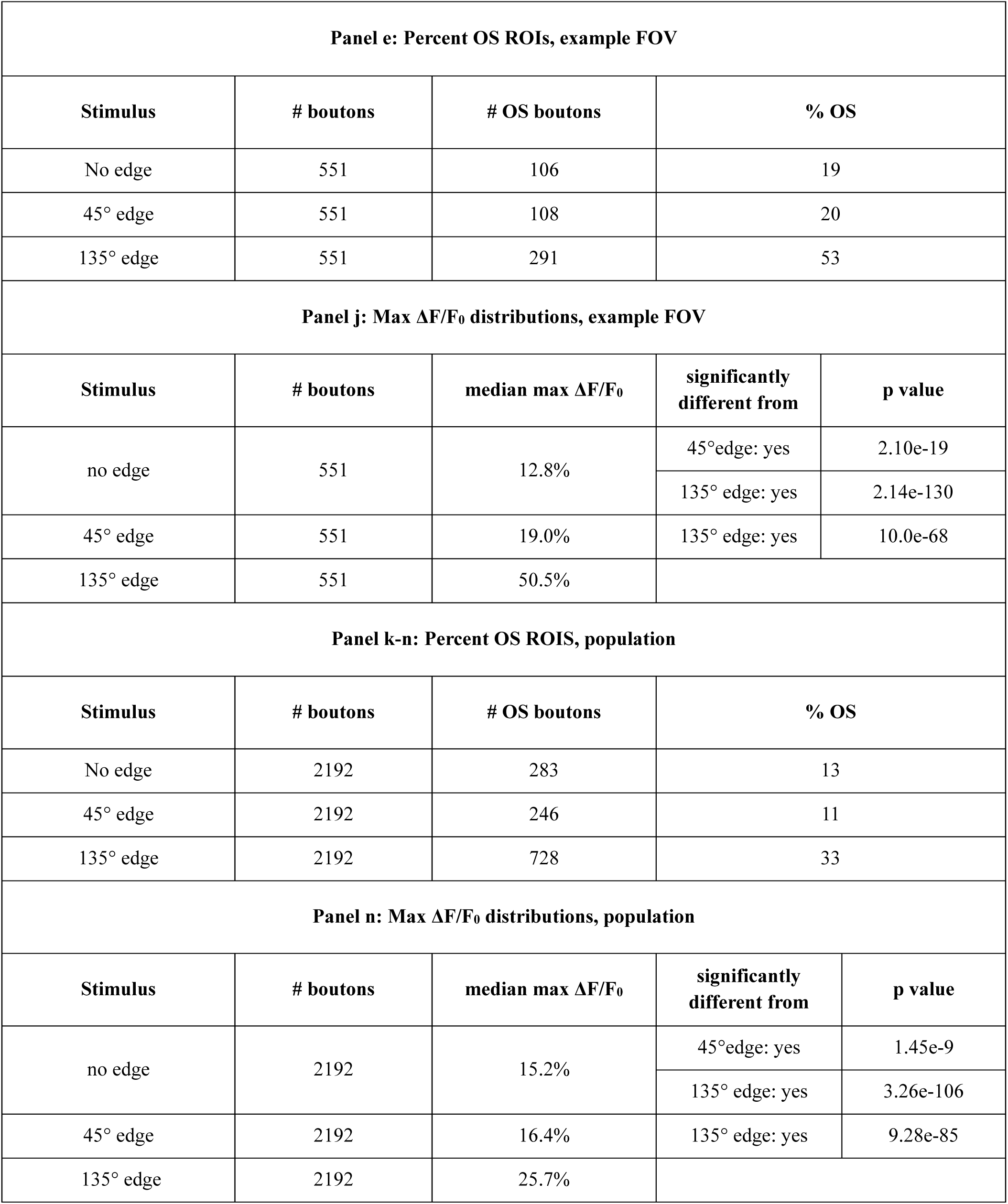
Percent OS ROIs and max ΔF/F_0_ distributions for Supp. Fig. 5.

**Supplementary Table 6:** Max ΔF/F_0_ and gOSI distributions for Fig 4.

| **Panel e: Max $\Delta F/F_0$ distributions, natively tuned boutons** |  |  |  |  |
| --- | --- | --- | --- | --- |
| **Stimulus** | **# boutons** | **median max $\Delta F/F_0$** | **significantly different from** | **p value** |
| no edge | 1128 | 30.5% | vertical edge: yes | 2.95e-19 |
|  |  |  | horizontal edge: yes | 1.30e-44 |
| vertical edge | 1128 | 41.9% | horizontal edge: yes | 7.83e-9 |
| horizontal edge | 1128 | 50.9% |  |  |
| **Panel g: gOSI distributions, natively tuned boutons** |  |  |  |  |
| no edge | 1128 | 0.38 | vertical edge: yes | 3.28e-17 |
|  |  |  | horizontal edge: yes | 3.61e-78 |
| vertical edge | 1082 | 0.29 | horizontal edge: yes | 5.23e-30 |
| horizontal edge | 1112 | 0.20 |  |  |
| **Panel k: Max $\Delta F/F_0$ distributions, non-natively tuned boutons** |  |  |  |  |
| **Stimulus** | **# boutons** | **median max $\Delta F/F_0$** | **significantly different from** | **p value** |
| no edge | 2892 | 14.6% | vertical edge: yes | 5.60e-186 |
|  |  |  | horizontal edge: yes | 0 |
| vertical edge | 2892 | 27.6% | horizontal edge: yes | 3.02e-33 |
| horizontal edge | 2892 | 36.4% |  |  |
| **Panel m: gOSI distributions, non-natively tuned boutons** |  |  |  |  |
| **Stimulus** | **# boutons** | **median gOSI** | **significantly different from** | **p value** |
| no edge | 2080 | 0.20 | vertical edge: yes | 1.18e-137 |
|  |  |  | horizontal edge: yes | 1.65e-84 |
| vertical edge | 2709 | 0.36 | horizontal edge: yes | 1.04e-10 |
| horizontal edge | 2830 | 0.32 |  |  |

**Supplementary Table 7:** Change in gOSI and max ΔF/F_0_ for non-natively tuned boutons for Supp. Fig. 6.

| Panel a: Change in gOSI, vertical and horizontal edges |  |  |  |  |  |  |  |
| --- | --- | --- | --- | --- | --- | --- | --- |
| OS condition | # boutons | Stimulus | median ΔgOSI | lower quartile | upper quartile | significantly different from | p value |
| Tuned to both edges | 838 | vertical edge | 0.17 | 0.05 | 0.30 | 0: yes | 7.80e-58 |
|  |  |  |  |  |  | H edge: yes | 3.20e-5 |
|  |  | horizontal edge | 0.12 | -0.02 | 0.27 | 0: yes | 2.09e-35 |
| Tuned to vertical edge only | 1003 | vertical edge | 0.16 | 0.04 | 0.28 | 0: yes | 1.80e-86 |
|  |  |  |  |  |  | H edge: yes | 5.90e-98 |
|  |  | horizontal edge | -0.05 | -0.18 | 0.06 | 0: yes | 9.70e-16 |
| Tuned to horizontal edge only | 1051 | vertical edge | -0.005 | -0.17 | 0.13 | 0: no | 0.138 |
|  |  |  |  |  |  | H edge: yes | 1.29e-37 |
|  |  | horizontal edge | 0.14 | -0.01 | 0.29 | 0: yes | 6.30e-49 |
| Not tuned to either edge | 2069 | vertical edge | -0.01 | -0.16 | 0.12 | 0: yes | 9.20e-5 |
|  |  |  |  |  |  | H edge: yes | 0.006 |
|  |  | horizontal edge | -0.03 | -0.18 | 0.10 | 0: yes | 7.97e-15 |
| Panel b: Change in max ΔF/F<sub>0</sub>, vertical and horizontal edges |  |  |  |  |  |  |  |
| OS condition | # boutons | Stimulus | median Δ(max ΔF/F<sub>0</sub>) | lower quartile | upper quartile | significantly different from | p value |
| Tuned to both edges | 838 | vertical edge | 18.6 | 9.0 | 32.5 | 0: yes | 7.56e-120 |
|  |  |  |  |  |  | H edge: yes | 2.63e-25 |
|  |  | horizontal edge | 29.2 | 15.5 | 50.4 | 0: yes | 3.87e-133 |
| Tuned to vertical edge only | 1003 | vertical edge | 18.4 | 8.5 | 35.5 | 0: yes | 8.53e-141 |
|  |  |  |  |  |  | H edge: yes | 2.94e-22 |
|  |  | horizontal edge | 10.5 | 2.8 | 23.4 | 0: yes | 2.08e-104 |
| Tuned to horizontal edge only | 1051 | vertical edge | 4.5 | -1.6 | 11.6 | 0: yes | 1.08e-46 |
|  |  |  |  |  |  | H edge: yes | 3.18e-137 |
|  |  | horizontal edge | 22.5 | 10.7 | 40.3 | 0: yes | 1.02e-160 |
| Not tuned to either edge | 2069 | vertical edge | 4.0 | -1.2 | 10.2 | 0: yes | 2.27e-92 |
|  |  |  |  |  |  | H edge: yes | 2.97e-7 |

|  |  | horizontal edge | 5.3 | -0.3 | 12.2 | 0: yes | 1.00e-131 |
| --- | --- | --- | --- | --- | --- | --- | --- |
| Panel c: Change in gOSI, 45° and 135° edges |  |  |  |  |  |  |  |
| OS condition | # boutons | Stimulus | median ΔgOSI | lower quartile | upper quartile | significantly different from | p value |
| both edges | 81 | 45° edge | 0.09 | -0.07 | 0.30 | 0: yes | 0.0017 |
|  |  |  |  |  |  | 135° edge: no | 0.2561 |
|  |  | 135° edge | 0.05 | -0.06 | 0.19 | 0: yes | 0.0246 |
| 45° edge only | 114 | 45° edge | 0.24 | 0.06 | 0.37 | 0: yes | 3.62e-12 |
|  |  |  |  |  |  | 135° edge: yes | 1.45e-15 |
|  |  | 135° edge | -0.04 | -0.22 | 0.09 | 0: yes | 0.0044 |
| 135° edge only | 536 | 45° edge | -0.04 | -0.17 | 0.08 | 0: yes | 4.25e-6 |
|  |  |  |  |  |  | 135° edge: yes | 3.31e-31 |
|  |  | 135° edge | 0.13 | -0.02 | 0.25 | 0: yes | 1.53e-28 |
| neither edge | 1145 | 45° edge | -0.02 | -0.17 | 0.13 | 0: yes | 2.48e-4 |
|  |  |  |  |  |  | 135° edge: no | 0.0910 |
|  |  | 135° edge | -0.02 | -0.19 | 0.11 | 0: yes | 4.51e-9 |
| Panel d: Change in max ΔF/F<sub>0</sub>, 45° and 135° edges |  |  |  |  |  |  |  |
| OS condition | # boutons | Stimulus | median Δmax ΔF/F<sub>0</sub> | lower quartile | upper quartile | significantly different from | p value |
| both edges | 81 | 45° edge | 17.9 | 6.8 | 37.1 | 0: yes | 3.68e-14 |
|  |  |  |  |  |  | 135° edge: | 2.49e-8 |
|  |  | 135° edge | 48.0 | 22.0 | 91.9 | 0: yes | 3.55e-15 |
| 45° edge only | 114 | 45° edge | 5.2 | -0.5 | 18.9 | 0: yes | 4.64e-10 |
|  |  |  |  |  |  | 135° edge: no | 0.36 |
|  |  | 135° edge | 8.3 | -0.8 | 25.3 | 0: yes | 6.90e-11 |
| 135° edge only | 536 | 45° edge | 2.5 | -3.7 | 9.3 | 0: yes | 3.06e-9 |
|  |  |  |  |  |  | 135° edge: yes | 4.65e-84 |
|  |  | 135° edge | 23.8 | 11.6 | 51.1 | 0: yes | 5.47e-81 |
| neither edge | 1145 | 45° edge | 1.0 | -4.8 | 6.0 | 0: yes | 0.003 |
|  |  |  |  |  |  | 135° edge: yes | 1.36e-22 |
|  |  | 135° edge | 4.2 | -1.9 | 11.8 | 0: yes | 1.52e-45 |

**Supplementary Table 8:** Max ΔF/F_0_ and gOSI distributions for Supp. Fig 7.

| **Panel e: Max $\Delta F/F_0$ distributions, natively tuned boutons** |  |  |  |  |
| --- | --- | --- | --- | --- |
| **Stimulus** | **# boutons** | **median max $\Delta F/F_0$** | **significantly different from** | **p value** |
| no edge | 283 | 30.0% | 45° edge: yes | 2.49e-8 |
|  |  |  | 135° edge: yes | 4.88e-4 |
| 45° edge | 283 | 20.7% | 135° edge: yes | 4.27e-20 |
| 135° edge | 283 | 34.9% |  |  |
| **Panel g: gOSI distributions, natively tuned boutons** |  |  |  |  |
| **Stimulus** | **# boutons** | **median gOSI** | **significantly different from** | **p value** |
| no edge | 283 | 0.52 | 45° edge: yes | 2.69e-30 |
|  |  |  | 135° edge: yes | 1.01e-39 |
| 45° edge | 265 | 0.27 | 135° edge: yes | 0.0255 |
| 135° edge | 273 | 0.28 |  |  |
| **Panel k: Max $\Delta F/F_0$ distributions, non-natively tuned boutons** |  |  |  |  |
| **Stimulus** | **# boutons** | **median max $\Delta F/F_0$** | **significantly different from** | **p value** |
| no edge | 731 | 13.3% | 45° edge: yes | 1.69e-19 |
|  |  |  | 135° edge: yes | 3.06e-136 |
| 45° edge | 731 | 18.1% | 135° edge: yes | 1.03e-76 |
| 135° edge | 731 | 40.4% |  |  |
| **Panel m: gOSI distributions, non-natively tuned boutons** |  |  |  |  |
| **Stimulus** | **# boutons** | **median gOSI** | **significantly different from** | **p value** |
| no edge | 541 | 0.26 | 45° edge: yes | 0.0298 |
|  |  |  | 135° edge: yes | 1.26e-34 |
| 45° edge | 656 | 0.29 | 135° edge: yes | 9.21e-26 |
| 135° edge | 726 | 0.39 |  |  |

**Supplementary Table 9:** Max ΔF/F_0_ and Percent OS ROIs for Fig. 5.

| **Panel j: Max $\Delta F/F_0$ distributions** |  |  |  |  |
| --- | --- | --- | --- | --- |
| **Stimulus** | **# boutons** | **median $\Delta F/F_0$** | **significantly different from** | **p value** |
| no edge | 3966 | 19.9% | light gray edge: yes | 2.99e-87 |
|  |  |  | dark gray edge: yes | 8.57e-187 |
|  |  |  | black edge: yes | 0 |
| light gray edge | 3966 | 27.6% | dark gray edge: yes | 4.98e-24 |
|  |  |  | black edge: yes | 8.34e-138 |
| dark gray edge | 3966 | 33.3% | black edge: yes | 2.90e-49 |
| black edge | 3966 | 45.5% |  |  |
| **Percent OS ROIs** |  |  |  |  |
| **Stimulus** | **# boutons** | **% OS** | **significantly different from** | **p value** |
| no edge | 3966 | 12 | light gray edge: yes | 0.02 |
|  |  |  | dark gray edge: yes | 0.02 |
|  |  |  | black edge: yes | 0.02 |
| light gray edge | 3966 | 41 | dark gray edge: no | 0.38 |
|  |  |  | black edge: no | 0.22 |
| dark gray edge | 3966 | 50 | black edge: no | 0.81 |
| black edge | 3966 | 48 |  |  |

**Supplementary Table 10:**
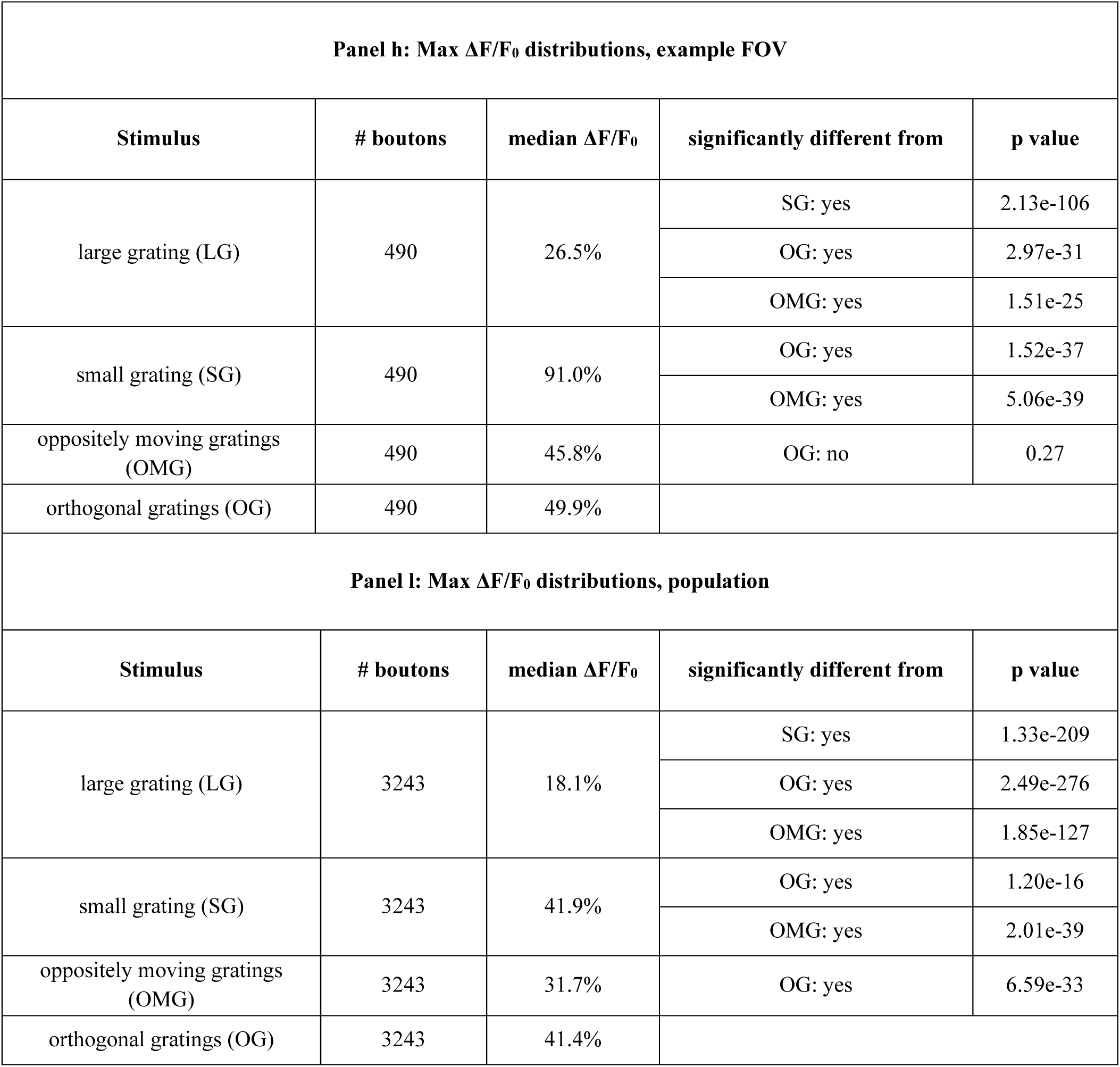
Max ΔF/F_0_ distributions for Fig. 6.

**Supplementary Table 11:** Max ΔF/F_0_ and gOSI distributions for Supp. Fig. 8.

| **Panel e: Max $\Delta F/F_0$ distributions, natively tuned boutons** |  |  |  |  |
| --- | --- | --- | --- | --- |
| **Stimulus** | **# boutons** | **median $\Delta F/F_0$** | **significantly different from** | **p value** |
| large grating (LG) | 738 | 32.2% | OG: yes | 1.31e-42 |
|  |  |  | OMG: yes | 4.27e-23 |
| orthogonal gratings (OG) | 738 | 57.5% | OMG: yes | 1.80e-5 |
| oppositely moving gratings (OMG) | 738 | 47.8% |  |  |
| **Panel g: gOSI distributions, natively tuned boutons** |  |  |  |  |
| **Stimulus** | **# boutons** | **median gOSI** | **significantly different from** | **p value** |
| large grating (LG) | 738 | 0.36 | OG: yes | 1.63e-24 |
|  |  |  | OMG: yes | 7.71e-25 |
| orthogonal gratings (OG) | 729 | 0.23 | OMG: no | 0.1454 |
| oppositely moving gratings (OMG) | 723 | 0.23 |  |  |
| **Panel k: Max $\Delta F/F_0$ distributions, non-natively tuned boutons** |  |  |  |  |
| **Stimulus** | **# boutons** | **median $\Delta F/F_0$** | **significantly different from** | **p value** |
| large grating (LG) | 1765 | 16.2% | OG: yes | 7.48e-271 |
|  |  |  | OMG: yes | 5.55e-143 |
| orthogonal gratings (OG) | 1765 | 45.8% | OMG: yes | 2.16e-34 |
| oppositely moving gratings (OMG) | 1765 | 33.0% |  |  |
| **Panel m: gOSI distributions, non-natively tuned boutons** |  |  |  |  |
| **Stimulus** | **# boutons** | **median gOSI** | **significantly different from** | **p value** |
| large grating (LG) | 1364 | 0.19 | OG: yes | 2.40e-68 |
|  |  |  | OMG: yes | 8.77e-56 |
| orthogonal gratings (OG) | 1742 | 0.30 | OMG: no | 0.0911 |
| oppositely moving gratings (OMG) | 1705 | 0.30 |  |  |

**Supplementary Table 12:** Change in gOSI and max ΔF/F_0_ for non-natively tuned boutons for Supp. Fig. 9.

| Panel a: Change in gOSI, example FOV |  |  |  |  |  |  |  |
| --- | --- | --- | --- | --- | --- | --- | --- |
| OS condition | # boutons | Stimulus | median ΔgOSI | lower quartile | upper quartile | significantly different from | p value |
| Tuned to both OMG and OG | 211 | OMG | 0.13 | 0.07 | 0.21 | 0: yes | 5.24e-36 |
|  |  |  |  |  |  | OG: no | 0.6618 |
|  |  | OG | 0.13 | 0.08 | 0.19 | 0: yes | 4.42e-36 |
| Tuned to OMG only | 47 | OMG | 0.11 | 0.07 | 0.16 | 0: yes | 4.56e-9 |
|  |  |  |  |  |  | OG: yes | 1.25e-7 |
|  |  | OG | 0.02 | -0.02 | 0.07 | 0: no | 0.0671 |
| Tuned to OG only | 46 | OMG | 0.01 | -0.04 | 0.07 | 0: no | 0.4675 |
|  |  |  |  |  |  | OG: yes | 1.10e-9 |
|  |  | OG | 0.12 | 0.07 | 0.19 | 0: yes | 8.81e-9 |
| Not tuned to OMG or OG | 54 | OMG | 0.01 | -0.04 | 0.07 | 0: no | 0.3242 |
|  |  |  |  |  |  | OG: no | 0.4406 |
|  |  | OG | -0.01 | -0.06 | 0.05 | 0: no | 0.8802 |
| Panel b: Change in max ΔF/F<sub>0</sub>, example FOV | | | | | | | |
| OS condition | # boutons | Stimulus | median Δmax ΔF/F<sub>0</sub> | lower quartile | upper quartile | significantly different from | p value |
| Tuned to both OMG and OG | 211 | OMG | 24.7 | 12.6 | 42.5 | 0: yes | 1.96e-36 |
|  |  |  |  |  |  | OG: no | 0.4378 |
|  |  | OG | 27.3 | 14.1 | 43.0 | 0: yes | 1.80e-36 |
| Tuned to OMG only | 47 | OMG | 18.4 | 7.9 | 28.9 | 0: yes | 1.41e-9 |
|  |  |  |  |  |  | OG: no | 0.1908 |
|  |  | OG | 13.7 | 7.4 | 27.2 | 0: yes | 2.51e-9 |
| Tuned to OG only | 46 | OMG | 7.1 | 2.0 | 10.9 | 0: yes | 2.61e-6 |
|  |  |  |  |  |  | OG: yes | 0.0253 |
|  |  | OG | 12.2 | 3.7 | 22.7 | 0: yes | 1.43e-8 |
| Not tuned to OMG or OG | 54 | OMG | 4.4 | -0.6 | 11.2 | 0: yes | 4.63e-5 |
|  |  |  |  |  |  | OG: no | 0.6036 |

|  |  | OG | 5.3 | 2.2 | 11.3 | 0: yes | 4.60e-7 |
| --- | --- | --- | --- | --- | --- | --- | --- |
| Panel c: Change in gOSI, population |  |  |  |  |  |  |  |
| OS condition | # boutons | Stimulus | median ΔgOSI | lower quartile | upper quartile | significantly different from | p value |
| Tuned to both<br>OMG and OG | 768 | OMG | 0.11 | 0.03 | 0.2 | 0: yes | 5.86e-60 |
|  |  |  |  |  |  | OG: no | 0.9300 |
|  |  | OG | 0.11 | 0.02 | 0.2 | 0: yes | 3.14e-56 |
| Tuned to OMG<br>only | 400 | OMG | 0.13 | 0.02 | 0.3 | 0: yes | 4.01e-29 |
|  |  |  |  |  |  | OG: yes | 2.88e-23 |
|  |  | OG | -0.01 | -0.1 | 0.1 | 0: no | 0.2769 |
| Tuned to OG<br>only | 597 | OMG | -0.01 | -0.1 | 0.1 | 0: no | 0.0847 |
|  |  |  |  |  |  | OG: yes | 1.55e-21 |
|  |  | OG | 0.1 | -0.02 | 0.2 | 0: yes | 2.38e-29 |
| Not tuned to<br>OMG or OG | 705 | OMG | -0.02 | -0.2 | 0.1 | 0: yes | 5.18e-4 |
|  |  |  |  |  |  | OG: no | 0.4322 |
|  |  | OG | -0.03 | -0.2 | 0.1 | 0: yes | 6.66e-6 |
| Panel d: Change in max ΔF/F<sub>0</sub>, population |  |  |  |  |  |  |  |
| OS condition | # boutons | Stimulus | median Δmax ΔF/F<sub>0</sub> | lower quartile | upper quartile | significantly different from | p value |
| Tuned to both<br>OMG and OG | 768 | OMG | 21.5 | 11.9 | 38.2 | 0: yes | 6.02e-124 |
|  |  |  |  |  |  | OG: yes | 3.96e-21 |
|  |  | OG | 33.7 | 18.1 | 57.5 | 0: yes | 3.04e-126 |
| Tuned to OMG<br>only | 400 | OMG | 17.8 | 8.6 | 30.1 | 0: yes | 1.06e-63 |
|  |  |  |  |  |  | OG: no | 0.0594 |
|  |  | OG | 15.2 | 5.7 | 29.8 | 0: yes | 1.44e-53 |
| Tuned to OG<br>only | 597 | OMG | 8.7 | 2.3 | 17.0 | 0: yes | 9.87e-57 |
|  |  |  |  |  |  | OG: yes | 2.48e-61 |
|  |  | OG | 24.9 | 13.4 | 41.7 | 0: yes | 6.87e-96 |
| Not tuned to<br>OMG or OG | 705 | OMG | 5.3 | -0.1 | 11.4 | 0: yes | 4.49e-47 |
|  |  |  |  |  |  | OG: yes | 6.64e-16 |
|  |  | OG | 9.6 | 3.0 | 18.3 | 0: yes | 3.04e-85 |

## References

1. Basso, M. A. & May, P. J. Circuits for Action and Cognition: A View from the Superior Colliculus. Annu. Rev. Vis. Sci. 3, 197–226 (2017).

2. Basso, M. A., Bickford, M. E. & Cang, J. Unraveling circuits of visual perception and cognition through the superior colliculus. Neuron 109, 918–937 (2021).

3. Cang, J., Savier, E., Barchini, J. & Liu, X. Visual Function, Organization, and Development of the Mouse Superior Colliculus. Annu. Rev. Vis. Sci. 4, 239–262 (2018).

4. Hoy, J. L. & Farrow, K. The superior colliculus. Curr. Biol. 35, R164–R168 (2025).

5. Krauzlis, R. J., Lovejoy, L. P. & Zénon, A. Superior Colliculus and Visual Spatial Attention. Annu. Rev. Neurosci. 36, 165–182 (2013).

6. Veale, R., Hafed, Z. M. & Yoshida, M. How is visual salience computed in the brain? Insights from behaviour, neurobiology and modelling. Philos. Trans. R. Soc. B Biol. Sci. 372, 20160113 (2017).

7. Wurtz, R. H. & Albano, J. E. Visual-Motor Function of the Primate Superior Colliculus. Annu. Rev. Neurosci. 3, 189– 226 (1980).

8. Ellis, E. M., Gauvain, G., Sivyer, B. & Murphy, G. J. Shared and distinct retinal input to the mouse superior colliculus and dorsal lateral geniculate nucleus. J. Neurophysiol. 116, 602–610 (2016).

9. Wang, Q. & Burkhalter, A. Stream-Related Preferences of Inputs to the Superior Colliculus from Areas of Dorsal and Ventral Streams of Mouse Visual Cortex. J. Neurosci. 33, 1696–1705 (2013).

10. Barchini, J., Shi, X., Chen, H. & Cang, J. Bidirectional encoding of motion contrast in the mouse superior colliculus. eLife 7, e35261 (2018).

11. Liang, Y., Lu, R., Borges, K. & Ji, N. Stimulus edges induce orientation tuning in superior colliculus. Nat. Commun. 14, 4756 (2023).

12. Yilmaz, M. & Meister, M. Rapid Innate Defensive Responses of Mice to Looming Visual Stimuli. Curr. Biol. 23, 2011– 2015 (2013).

13. Wu, R. et al. Preference-independent saliency map in the mouse superior colliculus. Commun. Biol. 8, 565 (2025).

14. Itti, L., Koch, C. & Niebur, E. A model of saliency-based visual attention for rapid scene analysis. IEEE Trans. Pattern Anal. Mach. Intell. 20, 1254–1259 (1998).

15. Itti, L. & Koch, C. Computational modelling of visual attention. Nat. Rev. Neurosci. 2, 194–203 (2001).

16. Yan, Y., Zhaoping, L. & Li, W. Bottom-up saliency and top-down learning in the primary visual cortex of monkeys. Proc. Natl. Acad. Sci. 115, 10499–10504 (2018).

17. Zhang, X., Zhaoping, L., Zhou, T. & Fang, F. Neural Activities in V1 Create a Bottom-Up Saliency Map. Neuron 73, 183–192 (2012).

18. Zhaoping, L. A saliency map in primary visual cortex. (2002).

19. Zhaoping, L. Parallel Advantage: Further Evidence for Bottom-up Saliency Computation by Human Primary Visual Cortex. Perception 51, 60–69 (2022).

20. White, B. J., Kan, J. Y., Levy, R., Itti, L. & Munoz, D. P. Superior colliculus encodes visual saliency before the primary visual cortex. Proc. Natl. Acad. Sci. 114, 9451–9456 (2017).

21. De Malmazet, D., Kühn, N. K., Li, C. & Farrow, K. Retinal origin of orientation but not direction selective maps in the superior colliculus. Curr. Biol. 34, 1222–1233.e7 (2024).

22. Molotkov, D., Ferrarese, L., Boissonnet, T. & Asari, H. Topographic axonal projection at single-cell precision supports local retinotopy in the mouse superior colliculus. Nat. Commun. 14, 7418 (2023).

23. Shi, X. et al. Retinal origin of direction selectivity in the superior colliculus. Nat. Neurosci. 20, 550–558 (2017).

24. Sibille, J. et al. High-density electrode recordings reveal strong and specific connections between retinal ganglion cells and midbrain neurons. Nat. Commun. 13, 5218 (2022).

25. Zhao, X., Liu, M. & Cang, J. Visual Cortex Modulates the Magnitude but Not the Selectivity of Looming-Evoked Responses in the Superior Colliculus of Awake Mice. Neuron 84, 202–213 (2014).

26. Ahmadlou, M., Tafreshiha, A. & Heimel, J. A. Visual Cortex Limits Pop-Out in the Superior Colliculus of Awake Mice. Cereb. Cortex 27, 5772–5783 (2017).

27. Liang, F. et al. Sensory Cortical Control of a Visually Induced Arrest Behavior via Corticotectal Projections. Neuron 86, 755–767 (2015).

28. Chen, T.-W. et al. Ultrasensitive fluorescent proteins for imaging neuronal activity. Nature 499, 295–300 (2013).

29. Ji, N., Milkie, D. E. & Betzig, E. Adaptive optics via pupil segmentation for high-resolution imaging in biological tissues. Nat. Methods 7, 141–147 (2010).

30. Zhang, Q. et al. Adaptive optics for optical microscopy [Invited]. Biomed. Opt. Express 14, 1732 (2023).

31. Pachitariu, M. et al. Suite2p: beyond 10,000 neurons with standard two-photon microscopy. Preprint at 10.1101/061507 (2016).

32. Baden, T. et al. The functional diversity of retinal ganglion cells in the mouse. Nature 529, 345–350 (2016).

33. Nath, A. & Schwartz, G. W. Cardinal Orientation Selectivity Is Represented by Two Distinct Ganglion Cell Types in Mouse Retina. J. Neurosci. 36, 3208–3221 (2016).

34. Hong, Y. K., Kim, I. & Sanes, J. R. Stereotyped axonal arbors of retinal ganglion cell subsets in the mouse superior colliculus. J. Comp. Neurol. 519, 1691–1711 (2011).

35. Rodieck, R. W. Quantitative analysis of cat retinal ganglion cell response to visual stimuli. Vision Res. 5, 583–601 (1965).

36. Cooler, S. & Schwartz, G. W. An offset ON–OFF receptive field is created by gap junctions between distinct types of retinal ganglion cells. Nat. Neurosci. 24, 105–115 (2021).

37. Krieger, B., Qiao, M., Rousso, D. L., Sanes, J. R. & Meister, M. Four alpha ganglion cell types in mouse retina: Function, structure, and molecular signatures. PLOS ONE 12, e0180091 (2017).

38. Marco, S. D., Protti, D. A. & Solomon, S. G. Excitatory and inhibitory contributions to receptive fields of alpha-like retinal ganglion cells in mouse. J. Neurophysiol. 110, 1426–1440 (2013).

39. Kuo, A., Gardner, J. L. & Merriam, E. P. Shared computational principles for mouse superior colliculus and primate population orientation selectivity. Preprint at 10.1101/2025.02.22.639626 (2025).

40. Girman, S. V. & Lund, R. D. Most Superficial Sublamina of Rat Superior Colliculus: Neuronal Response Properties and Correlates With Perceptual Figure–Ground Segregation. J. Neurophysiol. 98, 161–177 (2007).

41. Huang, X., Rangel, M., Briggman, K. L. & Wei, W. Neural mechanisms of contextual modulation in the retinal direction selective circuit. Nat. Commun. 10, 2431 (2019).

42. White, B. J. et al. Superior colliculus neurons encode a visual saliency map during free viewing of natural dynamic video. Nat. Commun. 8, 14263 (2017).

43. Koch, C. & Ullman, S. Shifts in Selective Visual Attention: Towards the Underlying Neural Circuitry. in Matters of Intelligence (ed. Vaina, L. M.) 115–141 (Springer Netherlands, Dordrecht, 1987). doi:10.1007/978-94-009-3833-5_5.

44. Nath, A. & Schwartz, G. W. Electrical synapses convey orientation selectivity in the mouse retina. Nat. Commun. 8, 2025 (2017).

45. Antinucci, P., Suleyman, O., Monfries, C. & Hindges, R. Neural Mechanisms Generating Orientation Selectivity in the Retina. Curr. Biol. 26, 1802–1815 (2016).

46. Antinucci, P. & Hindges, R. Orientation-Selective Retinal Circuits in Vertebrates. Front. Neural Circuits 12, 11 (2018).

47. Venkataramani, S. & Taylor, W. R. Orientation Selectivity in Rabbit Retinal Ganglion Cells Is Mediated by Presynaptic Inhibition. J. Neurosci. 30, 15664–15676 (2010).

48. Venkataramani, S. & Taylor, W. R. Synaptic Mechanisms Generating Orientation Selectivity in the ON Pathway of the Rabbit Retina. J. Neurosci. 36, 3336–3349 (2016).

49. Girman, S. & Lund, R. Orientation-Specific Modulation of Rat Retinal Ganglion Cell Responses and Its Dependence on Relative Orientations of the Center and Surround Gratings. J. Neurophysiol. 104, 2951–2962 (2010).

50. Kuffler, S. W. DISCHARGE PATTERNS AND FUNCTIONAL ORGANIZATION OF MAMMALIAN RETINA. J. Neurophysiol. 16, 37–68 (1953).

51. Chiao, C.-C. & Masland, R. H. Contextual tuning of direction-selective retinal ganglion cells. Nat. Neurosci. 6, 1251– 1252 (2003).

52. Baccus, S. A., Olveczky, B. P., Manu, M. & Meister, M. A Retinal Circuit That Computes Object Motion. J. Neurosci. 28, 6807–6817 (2008).

53. Ölveczky, B. P., Baccus, S. A. & Meister, M. Segregation of object and background motion in the retina. Nature 423, 401–408 (2003).

54. Rivlin-Etzion, M., Wei, W. & Feller, M. B. Visual Stimulation Reverses the Directional Preference of Direction-Selective Retinal Ganglion Cells. Neuron 76, 518–525 (2012).

55. Cazemier, J. L. et al. Involvement of superior colliculus in complex figure detection of mice. eLife 13, e83708 (2024).

56. Patterson, S. S. et al. An S-cone circuit for edge detection in the primate retina. Sci. Rep. 9, 11913 (2019).

57. Van Wyk, M., Taylor, W. R. & Vaney, D. I. Local Edge Detectors: A Substrate for Fine Spatial Vision at Low Temporal Frequencies in Rabbit Retina. J. Neurosci. 26, 13250–13263 (2006).

58. Gaynes, J. A., Budoff, S. A., Grybko, M. J., Hunt, J. B. & Poleg-Polsky, A. Classical center-surround receptive fields facilitate novel object detection in retinal bipolar cells. Nat. Commun. 13, 5575 (2022).

59. Gale, S. D. & Murphy, G. J. Distinct Representation and Distribution of Visual Information by Specific Cell Types in Mouse Superficial Superior Colliculus. J. Neurosci. 34, 13458–13471 (2014).

60. Gale, S. D. & Murphy, G. J. Active Dendritic Properties and Local Inhibitory Input Enable Selectivity for Object Motion in Mouse Superior Colliculus Neurons. J. Neurosci. 36, 9111–9123 (2016).

61. Gale, S. D. & Murphy, G. J. Distinct cell types in the superficial superior colliculus project to the dorsal lateral geniculate and lateral posterior thalamic nuclei. J. Neurophysiol. 120, 1286–1292 (2018).

62. Li, Y. & Meister, M. Functional cell types in the mouse superior colliculus. eLife 12, e82367 (2023).

63. Cui, P. et al. Recurrent circuits encode de novo visual center-surround computations in the mouse superior colliculus. PLOS Biol. 23, e3003414 (2025).

64. Binns, K. E. & Salt, T. E. Different roles for GABA_A_ and GABA_B_ receptors in visual processing in the rat superior colliculus. J. Physiol. 504, 629–639 (1997).

65. Kaneda, K. & Isa, T. GABAergic mechanisms for shaping transient visual responses in the mouse superior colliculus. Neuroscience 235, 129–140 (2013).

66. Liu, Y. et al. Mapping visual functions onto molecular cell types in the mouse superior colliculus. Neuron 111, 1876–1886.e5 (2023).

67. Zhaoping, L. From the optic tectum to the primary visual cortex: migration through evolution of the saliency map for exogenous attentional guidance. Curr. Opin. Neurobiol. 40, 94–102 (2016).

68. Knudsen, E. I. Control from below: the role of a midbrain network in spatial attention. Eur. J. Neurosci. 33, 1961–1972 (2011).

69. Knudsen, E. I., Schwarz, J. S., Knudsen, P. F. & Sridharan, D. Space-Specific Deficits in Visual Orientation Discrimination Caused by Lesions in the Midbrain Stimulus Selection Network. Curr. Biol. 27, 2053–2064.e5 (2017).

70. Lintz, M. J., Essig, J., Zylberberg, J. & Felsen, G. Spatial representations in the superior colliculus are modulated by competition among targets. Neuroscience 408, 191–203 (2019).

71. Wolf, A. B. et al. An integrative role for the superior colliculus in selecting targets for movements. J. Neurophysiol. 114, 2118–2131 (2015).

72. Binns, K. E. & Salt, T. E. The functional influence of nicotinic cholinergic receptors on the visual responses of neurones in the superficial superior colliculus. Vis. Neurosci. 17, 283–289 (2000).

73. Chen, C. & Regehr, W. G. Presynaptic Modulation of the Retinogeniculate Synapse. J. Neurosci. 23, 3130–3135 (2003).

74. Govindaiah, G. & Cox, C. L. Depression of retinogeniculate synaptic transmission by presynaptic D_2_-like dopamine receptors in rat lateral geniculate nucleus. Eur. J. Neurosci. 23, 423–434 (2006).

75. Miller, R. J. Presynaptic receptors. Annu. Rev. Pharmacol. Toxicol. 38, 201–227 (1998).

76. Mooney, R. D., Shi, M. Y. & Rhoades, R. W. Modulation of retinotectal transmission by presynaptic 5-HT1B receptors in the superior colliculus of the adult hamster. J. Neurophysiol. 72, 3–13 (1994).

77. Prusky, G. T. & Cynader, M. S. [^3^ H]nicotine binding sites are associated with mammalian optic nerve terminals. Vis. Neurosci. 1, 245–248 (1988).

78. Reggiani, J. D. S. et al. Brainstem serotonin neurons selectively gate retinal information flow to thalamus. Neuron 111, 711–726.e11 (2023).

79. Seeburg, D. P., Liu, X. & Chen, C. Frequency-Dependent Modulation of Retinogeniculate Transmission by Serotonin. J. Neurosci. 24, 10950–10962 (2004).

80. Beltramo, R. & Scanziani, M. A collicular visual cortex: Neocortical space for an ancient midbrain visual structure. Science 363, 64–69 (2019).

81. Brenner, J. M., Beltramo, R., Gerfen, C. R., Ruediger, S. & Scanziani, M. A genetically defined tecto-thalamic pathway drives a system of superior-colliculus-dependent visual cortices. Neuron 111, 2247–2257.e7 (2023).

82. Cone, J. J., Mitchell, A. O., Parker, R. K. & Maunsell, J. H. R. Stimulus-dependent differences in cortical versus subcortical contributions to visual detection in mice. Curr. Biol. 34, 1940–1952.e5 (2024).

83. Hoy, J. L., Bishop, H. I. & Niell, C. M. Defined Cell Types in Superior Colliculus Make Distinct Contributions to Prey Capture Behavior in the Mouse. Curr. Biol. 29, 4130–4138.e5 (2019).

84. Ito, S. & Feldheim, D. A. The Mouse Superior Colliculus: An Emerging Model for Studying Circuit Formation and Function. Front. Neural Circuits 12, 10 (2018).

85. Wang, L., McAlonan, K., Goldstein, S., Gerfen, C. R. & Krauzlis, R. J. A Causal Role for Mouse Superior Colliculus in Visual Perceptual Decision-Making. J. Neurosci. 40, 3768–3782 (2020).

86. Kühn, N. K. et al. Dendritic Architecture Enables de Novo Computation of Salient Motion in the Superior Colliculus. Preprint at 10.1101/2025.02.28.640764 (2025).

87. Dhande, O. S. & Huberman, A. D. Retinal ganglion cell maps in the brain: implications for visual processing. Curr. Opin. Neurobiol. 24, 133–142 (2014).

88. Zhaoping, L. A new framework for understanding vision from the perspective of the primary visual cortex. Curr. Opin. Neurobiol. 58, 1–10 (2019).

89. Yoshida, M. et al. Residual Attention Guidance in Blindsight Monkeys Watching Complex Natural Scenes. Curr. Biol. 22, 1429–1434 (2012).

90. Croner, L. J. & Kaplan, E. Receptive fields of P and M ganglion cells across the primate retina. Vision Res. 35, 7–24 (1995).

91. Field, G. D. et al. Spatial Properties and Functional Organization of Small Bistratified Ganglion Cells in Primate Retina. J. Neurosci. 27, 13261–13272 (2007).

92. Gauthier, J. L. et al. Receptive Fields in Primate Retina Are Coordinated to Sample Visual Space More Uniformly. PLoS Biol. 7, e1000063 (2009).

93. Broussard, G. J. et al. In vivo measurement of afferent activity with axon-specific calcium imaging. Nat. Neurosci. 21, 1272–1280 (2018).

94. Ji, N., Sato, T. R. & Betzig, E. Characterization and adaptive optical correction of aberrations during in vivo imaging in the mouse cortex. Proc. Natl. Acad. Sci. 109, 22–27 (2012).

95. Brainard, D. H. The Psychophysics Toolbox. Spat. Vis. 10, 433–436 (1997).

96. Schindelin, J. et al. Fiji: an open-source platform for biological-image analysis. Nat. Methods 9, 676–682 (2012).

97. Guizar-Sicairos, M., Thurman, S. T. & Fienup, J. R. Efficient subpixel image registration algorithms. Opt. Lett. 33, 156 (2008).

98. Sun, W., Tan, Z., Mensh, B. D. & Ji, N. Thalamus provides layer 4 of primary visual cortex with orientation- and direction-tuned inputs. Nat. Neurosci. 19, 308–315 (2016).

